# Mechanisms of transcription attenuation and condensation of RNA polymerase II by RECQ5

**DOI:** 10.1101/2025.02.05.636647

**Authors:** M. Sebesta, K. Skubnik, W.S. Morton, M. Kravec, K. Linhartova, V. Klapstova, J. Novacek, K. Kubicek, V. Bryja, R. Vacha, R. Stefl

## Abstract

The elongation rates of RNA polymerase II (RNAPII) require precise control to prevent transcriptional stress, which can impede co-transcriptional pre-mRNA processing and contribute to many age- or disease-associated molecular changes (*e.g.*, loss of proteostasis)^1–5^. Additionally mesoscale organization of transcription is thought to control the transcriptional rates^6^ and multiple factors have been reported to form biomolecular condensates and integrate RNAPII through the interaction with the C-terminal domain (CTD) of the largest subunit, RPB1^7–10^. However, the structural organization of these condensates remains uncharacterized due to their small size and inherently dynamic nature. Here, we investigated the molecular mechanisms by which a general transcription factor – RECQ5 – associates with hyperphosphorylated RNAPII elongation complex (P-RNAPII EC) and controls translocation of RNAPII along genes. We combined biochemical reconstitution, electron cryomicroscopy, cryotomography, and coarse-grained simulations. We report two mechanisms by which RECQ5 modulates RNAPII transcription. At the atomic level, we demonstrate that RECQ5 uses the brake helix as a doorstop to control RNAPII translocation along DNA, attenuating transcription. At the mesoscale level, RECQ5 forms a condensate scaffold matrix, integrating P-RNAPII EC through a network of site-specific interactions, reinforcing the condensate’s structural integrity. Our integrative, multi-scale study provides insights into the structural basis of transcription attenuation and into the molecular architecture and biogenesis of a model RNAPII condensate.

## Main Text

A model of condensate-based transcriptional organization has emerged from recent studies^11^, suggesting that the transcribing enzyme – RNA polymerase II (RNAPII) – may cluster into condensates associated with transcription initiation and elongation^7–10^. Transcriptional condensates bring together the necessary components to facilitate efficient and coordinated spatiotemporal activation and regulation of transcription^12^, and their improper biogenesis has been linked to disease^13,14^. RNAPII is recruited into these condensates through its low-complexity, C-terminal domain (CTD)^15^. Shuttling of RNAPII between the condensates depends on the phosphorylation code of the CTD^8,16,17^, in coordination with the transcriptional cycle^18^. Super-resolution and light-sheet imaging at submicron resolution have provided only limited insight into these condensates, due to inherent limits of their resolution, and their architecture remains uncharted^19,20^. Additionally, there is still limited understanding of whether the formation of RNAPII condensates is primarily driven by non-specific or site-specific, multivalent interactions^21,22^.

Transcriptional stress, caused by accelerated transcription elongation and elevated stalling of RNAPII, contributes to the age-associated decline of cellular and organismal homeostasis^1,2^. RNAPII elongation is controlled by a general elongation factor RECQ5, which reduces the elongation rate while decreasing pausing and arrest events to smooth transcription and prevent transcriptional stress^23^. This attenuation of RNAPII transcription is independent of the ATPase activity of RECQ5^24^, but requires binding of RECQ5 to the elongating RNAPII, which is poorly understood^25–28^. Previous studies with truncated RECQ5 variants reported that the interaction with RNAPII is mediated by the RPB1 subunit of RNAPII^25^, through its serine 2 and 5 phosphorylated CTD^29^ and jaw domain^30^ with Set2-Rpb1–interacting (SRI) and internal RNAPII–interacting (IRI) domains of RECQ5, respectively. Yet, the direct interaction of the full-length RECQ5 and elongating RNAPII has not been characterized biochemically nor structurally. We investigated the interaction between RNAPII elongation complex (RNAPII EC) and RECQ5 at different scales using a combination of electron cryomicroscopy (cryo-EM), cryotomography (cryo-ET), and molecular dynamics (MD) simulations. The collective structural insights outline a mechanistic model of transcription attenuation via RECQ5 recruitment and provide strong support for the condensate-based model of transcriptional organization^9,12,31^.

### The brake helix of RECQ5 IRI domain contacts downstream DNA in RNAPII EC and inhibits transcription

To elucidate the molecular mechanism governing the attenuation of RNAPII transcription by RECQ5, we initially tested the role of the two RNAPII interacting domains of RECQ5. We used their binding-deficient variants in the IRI (RECQ5^K58R, L556E, L602E^)^30^ and SRI (RECQ5^K58R, R943A^)^29^ domains, respectively, in an *in vitro* transcription assay using the human histone H3.3 template^30,32,33^ and phosphorylated RNAPII (P-RNAPII; Supplementary Fig. 1). To rule out possible interference caused by ATP depletion by RECQ5 translocation activity, the tested variants harbored substitution in the Walker A motif of the ATPase core (K58R), as established previously^24,30^. Firstly, the RECQ5 variants exhibited the expected binding profile towards isolated RPB1 domains (RECQ5^K58R, L556E, L602E^ not recognizing the jaw domain, whilst the RECQ5^K58R, R943A^ not recognizing the phosphorylated CTD) (Supplementary Fig. 1)^29,30^. RECQ5^K58R^ effectively inhibited RNAPII activity *in vitro* (Fig. 1a and Extended Data Fig. 1), thereby re-affirming that its helicase activity is not necessary for the repression. We observed that the RECQ5^K58R, R943A^ (SRI) variant efficiently inhibited transcription, whilst the RECQ5^K58R, L556E, L602E^ (IRI) variant did not. This suggests that the RECQ5 interaction with the phosphorylated CTD (P-CTD) of RNAPII is dispensable for the inhibition of RNAPII *in vitro*, whilst the interaction with the RPB1 jaw domain is essential (Fig. 1a).

**Fig. 1:**
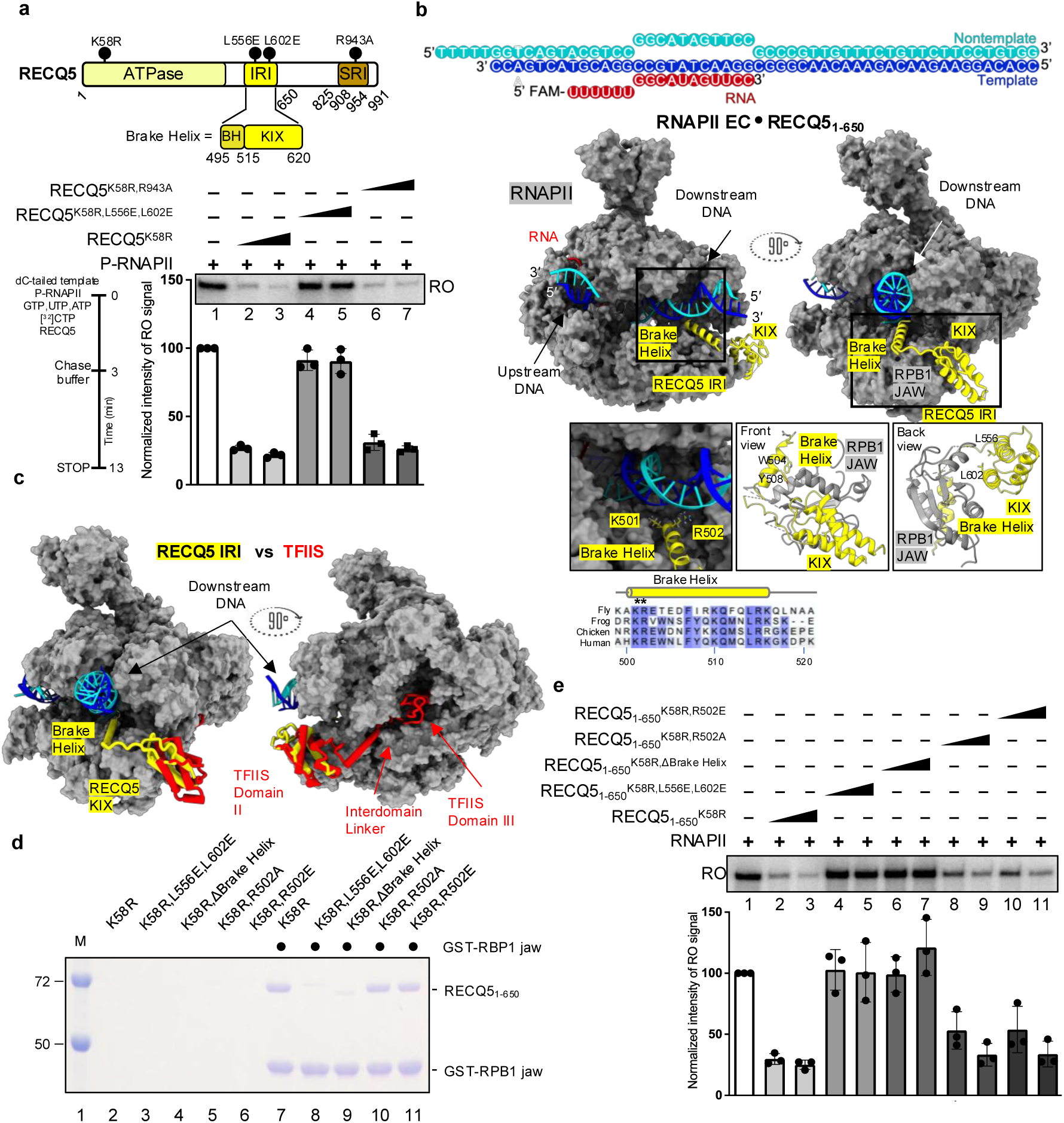
Domain architecture, cryo-EM structure of RNAPII EC•RECQ5_1-650_ complex, and transcription assays. **a**, Primary domain architecture of RECQ5 (upper). IRI domain but not SRI domain is required for transcription repression. Schematic overview of the *in vitro* transcription assay (lower). Run-off transcripts generated by phosphorylated RNAPII (P-RNAPII) in the presence of the indicated RECQ5 variants resolved by electrophoresis on a denaturing polyacrylamide gel and visualized by phosphorimaging. Quantified production of the run-off transcripts (RO). Error bars denote SD (n = 3). **b**, Nucleic acid scaffold used for RNAPII EC assembly (upper). Cartoon models of the three-dimensional cryo-EM structure view from the DNA entry side (middle). The 12-subunit P-RNAPII is shown in grey. RECQ5 is shown in yellow. DNA template and non-template strands are shown in blue and cyan, respectively. RNA is in red. The brake helix and the KIX domain interactions with RNAPII jaw domain and downstream DNA (lower). Sequence alignment of the brake helix. **c**, Superimposition of RNAPII complexes bound to RECQ5 and TFIIS (PDB:8A40). Color scheme as in b; TFIIS in red. **d**, Pull-down assay of RECQ5 variants with RPB1 jaw. **e**, The brake helix and R502 are required for transcription repression. Transcription assay as in a, performed with RNAPII.

The results from the transcription assays allowed us to use a C-terminal truncation of RECQ5 (RECQ5_1-650_) to reconstitute the RNAPII EC•RECQ5_1-650_ complex using a nucleic acid scaffold with DNA blunt ends (Fig. 1b), as overhangs could serve as artificial binding sites for RECQ5 helicase^34^. We imaged the complex by cryo-EM and resolved its structure to a nominal resolution of 3.2 Å (Supplementary Table 1 and Extended Data Figs. 2 and 3). The structure shows that the KIX domain, the C-terminal portion of the IRI domain, of RECQ5 binds the RPB1 jaw domain (Fig. 1b) at the site that partially overlaps with the known binding site for the elongation factor TFIIS (Fig. 1c)^35–38^, in agreement with a previous report^30^. However, our structure unveiled the N-terminal helix of the IRI domain, which we term the “brake helix”. This helix protrudes between the jaws of RNAPII and contacts the downstream DNA entering the RNAPII active center (Fig. 1b,c). Notably, our structure pinpointed the conserved arginine and lysine residues of the brake helix (K501 and R502), which interact with the sugar-phosphate backbone of both the template and non-template strands across the minor groove of the downstream DNA (Fig. 1b). Substituting the arginine residue with either alanine or glutamate, RECQ5^R502A^ or RECQ5^R502E^, had no impact on the RPB1 jaw binding, as demonstrated by *in vitro* pull-down assay (Fig. 1d). However, these substitutions resulted in impaired transcription repression (Fig. 1e). These findings affirm that the loss of inhibition is not a consequence of impaired binding to the RPB1 jaw, rather, it is likely attributed to the absence of contact between the arginine and the downstream DNA. To further investigate this repression-deficient variant, we determined 3.4-Å-resolution cryo-EM structure of RNAPII EC•RECQ5_1-650_^R502E^ complex (Supplementary Table 1 and Extended Data Fig. 4). The structure showed that the entire IRI domain of the RECQ5_1-650_^R502E^ maintains its interaction with the RPB1 jaw, analogous to the wild type, with the brake helix positioned at the DNA entry site. However, a weak density of the downstream DNA, relative to both the unbound and RECQ5_1-650_- associated RNAPII EC, suggests that the R502E charge-swapping substitution induces an electrostatic repulsion with the DNA (Extended Data Fig. 3d). The loss of DNA contacts renders the brake helix variant ineffective in stabilizing and controlling the passage of DNA through RNAPII.

Comparison of our structure with the TFIIS-bound RNAPII^38^ shows that TFIIS accesses the catalytic site through the bottom funnel side, whereas the brake helix of RECQ5 extends directly to the RPB1 jaw to contact downstream DNA from the front side (Fig. 1c). The partial overlap of the RNAPII binding sites for TFIIS and RECQ5 is consistent with the previous functional data^30^, which suggests that their binding to RPB1 jaw is mutually exclusive, requiring their exchange to control elongation under different circumstances (*e.g.*, to resume transcription of a stalled RNAPII EC complex). Importantly, the binding of RECQ5 to the RPB1 jaw of RNAPII does not interfere with other factors involved in elongation (Extended Data Fig. 3c). As RECQ5 is the only known factor contacting downstream DNA during elongation, we speculate that it might play a role during the encounter of RNAPII with a nucleosome.

### RECQ5 forms condensates that integrate RNAPII by site-specific interactions

To capture the interaction of P-CTD of RNAPII with RECQ5, we imaged the full-length P-RNAPII EC•RECQ5 complex using cryo-EM. Unexpectedly, cryo-EM imaging uncovered that the P-RNAPII EC•RECQ5 complex formed large, spherical objects, resembling condensates (Fig. 2a, left). Correlative light electron microscopy (CLEM) showed that these objects contained both RNAPII EC (labeled with fluorescein on the nucleic acid scaffold) and RECQ5 (fused with fluorescent protein – mCerulean) (Fig. 2b). Upon removal of a 200-amino acid, intrinsically disordered region (625-825; IDR) of RECQ5 (RNAPII EC•RECQ5^ΔIDR^; Fig. 2a, right), the large spherical objects were no longer visible on the cryo-EM grids. Single particle analysis (SPA) of these isolated RNAPII EC•RECQ5^ΔIDR^ particles provided a well-resolved map (at a resolution of 3.8 Å; Fig. 2a, right), which matched that of the RNAPII EC•RECQ5_1-650_ complex. However, no additional density corresponding to the SRI-CTD interactions was observed, even though the interaction between the SRI of RECQ5 and P-CTD was retained in the RECQ5^ΔIDR^ variant (Supplementary Fig. 2a). Collectively, these data suggest that RECQ5 may form condensates with liquid-like properties *in vitro*.

**Fig. 2:**
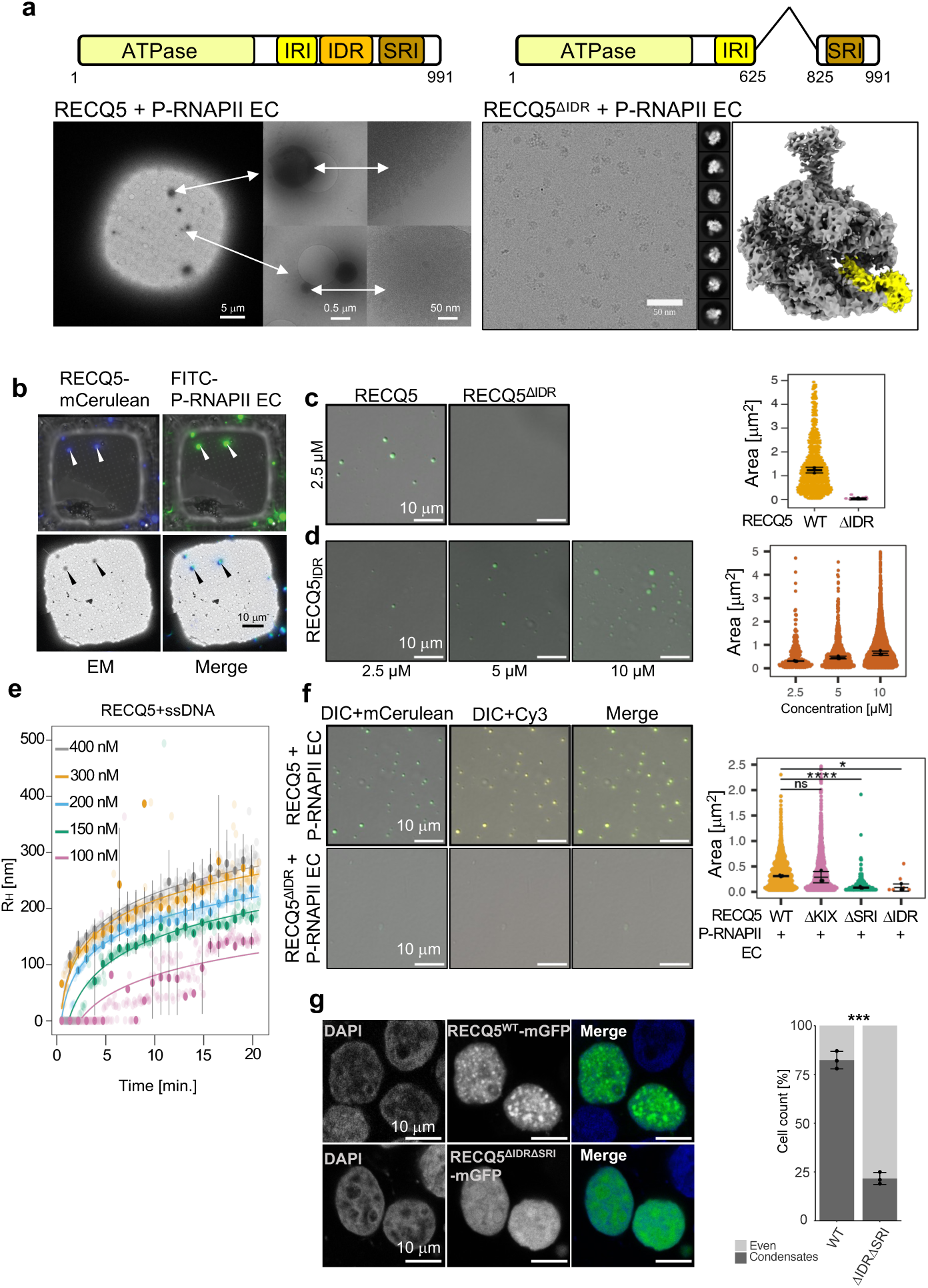
RECQ5 forms condensates and integrates P-RNAPII EC. **a**, Schematic domain overview of RECQ5, cryo-EM imaging of the P-RNAPII EC•RECQ5 and P-RNAPII EC•RECQ5^ΔIDR^ complexes (P-RNAPII at 1µM; RECQ5 variants at 5µM), 2D classes and reconstructed 3D cryo-EM map for P-RNAPII EC•RECQ5^ΔIDR^ complex. Region corresponding to RECQ5 is highlighted in yellow. **b**, Correlative light-electron microscopy (CLEM) of RECQ5- mCerulean and FITC-labeled P-RNAPII EC; DAPI channel for mCerulean and GFP channel for FITC. *In vitro* LLPS assay for mCerulean-labeled RECQ5 and RECQ5^ΔIDR^ (**c**) and isolated RECQ5_IDR_ (**d**). Representative images and quantifications (n=3), representing mean + SD. **e**, DLS data, hydrodynamic radius (nm), n=3, mean+SD. **f**, LLPS assay of Cy3-labeled P-RNAPII EC with RECQ5 and RECQ5^ΔIDR^, respectively. Statistical significance was determined by unpaired t-test. **g**, Ectopically expressed RECQ5-mGFP forms condensates in HEK293T cells, which requires both IDR and SRI domains. Micrographs of cells expressing RECQ5-mGFP (upper panel) and RECQ5^ΔIDRΔSRI^-mGFP (lower panel), and merge of both channels, stained for DAPI (nuclei). Right bar chart represents proportion of cells in which RECQ5-mGFP variants formed condensates (100 cells per experiment; n=3), mean+SD. Statistical significance was determined by unpaired t-test.

To investigate this phenomenon further, we used RECQ5-mCerulean and performed thorough biochemical characterization. We found that RECQ5-mCerulean, but not mCerulean alone, formed spherical condensates in a protein concentration-dependent manner in the presence of nucleic acid and/or crowding agent (polyethylene glycol; PEG) and these objects were sensitive to hexane-1,6-diol (Supplementary Fig. 2b-e). The key region responsible for the RECQ5 condensate formation *in vitro* was pinpointed by mapping with RECQ5 internal deletion variants tagged with monomeric mCerulean (RECQ5s-mCerulean; Fig. 2c,d and Supplementary Fig. 3). The mapping revealed that the deletion of the IDR domain (RECQ5^ΔIDR^) suppressed condensation of RECQ5, whilst the 200-amino acid fragment RECQ5_IDR_ in isolation did form condensates with characteristics consistent with liquid-liquid phase-separation (LLPS; Fig. 2 and Supplementary Fig. 2). Additionally, the increase in size of particles as determined by dynamic light scattering (DLS) indicated that RECQ5 undergoes phase-separation in the presence of ssDNA (Fig. 2e and Supplementary Fig. 4) at concentration as low as 0.1 μM. Subsequently, we tested the role of RECQ5 domains for heterotypic droplet formation with P-RNAPII EC and its components (Fig. 2f and Supplementary Figs. 5-7). We found that not only the deletion of the IDR domain (RECQ5^ΔIDR^) but also the deletion of the SRI domain (RECQ5^ΔSRI^) impaired heterotypic droplet formation with P-RNAPII EC, indicating that the specific binding of the SRI domain of RECQ5 to P-CTD of RPB1 is essential for heterotypic droplet formation. Likewise, removal of both RECQ5^ΔIDR^ and RECQ5^ΔSRI^ suppressed the formation of heterotypic droplets with P-CTD of RNAPII (Supplementary Fig. 8).

Fittingly, *in vivo*, the ectopic expression of RECQ5-mGFP in HEK293T cells led to the formation of condensates (Extended Data Fig. 5a), which was significantly diminished when cells expressed RECQ5^ΔIDRΔSRI^-mGFP variant (Fig. 2g). These data suggest that *in vivo*, both the interaction between RECQ5 and P-RNAPII, and the ability of RECQ5 to form condensates with liquid-like properties, are required for its condensation, as single deletion variants did retain wild-type levels of condensation (Extended Data Fig. 5b,c). This implies that RECQ5 might be associated with clustering to other condensates unrelated to transcription, reflecting its multi-faceted role *in vivo*^39^. Moreover, in a subset of condensates, RECQ5 colocalizes with RNAPII phosphorylated on S5 of the CTD (S5P-RNAPII) (Supplementary Fig. 9), suggesting that also *in vivo*, RECQ5 may contribute to the organization of S5P-RNAPII into condensates.

We conclude that RECQ5 forms condensates primarily through its IDR located within amino acids 625-825 (RECQ5_IDR_) and that the heterotypic condensates are further supported by site-specific interactions of the SRI domain of RECQ5 with the P-CTD of RPB1^38,39^. The RECQ5_IDR_ has a relatively uniform spacing of charged blocks along its primary sequence (Supplementary Fig. 3d), which are commonly thought to drive phase-separation^40–43^.

### Spatial organization of P-RNAPII EC•RECQ5 condensates

To reveal the spatial organization of P-RNAPII EC•RECQ5 condensates at molecular level, we reconstituted hyperphosphorylated RNAPII EC with RECQ5 at low, submicromolar concentrations and imaged it using cryo-EM and cryo-ET (Fig. 3a,b). The comparison of cryo-EM images of P-RNAPII EC alone and mixed with RECQ5 showed evenly dispersed P-RNAPII EC particles in their apo form, and the formation of ∼100 nm-sized condensates when P-RNAPII EC was bound with RECQ5 (Fig. 3b). On the cryo-EM grids with the P-RNAPII EC•RECQ5 complex, we also observed isolated RNAPII-type particles. SPA of the particles outside of the condensates revealed two classes of P-RNAPII EC: one bound to RECQ5 (∼2/3 of the particles) and the other in apo form (the remaining 1/3). The map reconstructed from the latter class displayed a typical RNAPII EC architecture (Fig 3b), which closely matched the previously determined structure^44^. Combined with the findings from CLEM (Fig. 2b), these observations suggest that RECQ5 predominantly resides within the condensates.

**Fig. 3:**
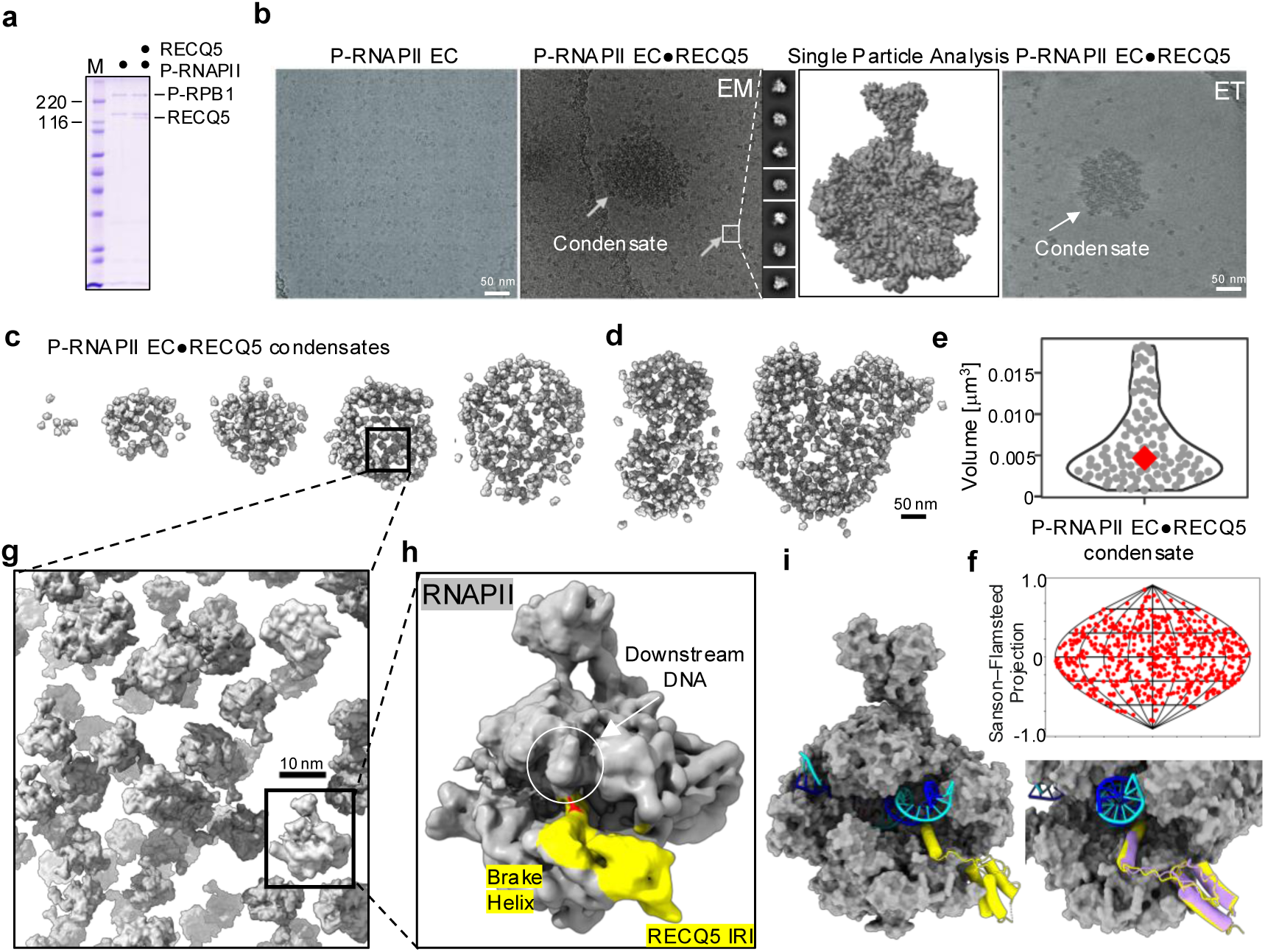
Cryo-ET of P-RNAPII EC•RECQ5 condensates. **a**, A scan of SDS-PAGE analysis of P-RNAPII and its complex with RECQ5 used for cryo-ET (P-RNAPII at 0.1 μM and RECQ5 at 0.25 μM). **b**, (from left to right) Cryo-EM micrographs of P-RNAPII EC and its complex with RECQ5. Example of 2D class averages and single particle cryo-EM reconstructions for off-condensate particles. Representative cryo-ET plane of the P-RNAPII EC•RECQ5 condensate. **c**, Representative final denoised 3D maps of selected condensates captured by cryo-ET analysis, ordered by increasing size. **d**, 3D cryo-ET maps of condensate coalescence. **e**, Volume analysis of the cryo-ET reconstructed P-RNAPII EC•RECQ5 condensates. **f**, Sinusoidal projection showing random P-RNAPII orientations in the condensate displayed in (d, left). **g**, The map from subtomogram averaging mapped back into the tomographic volume. **h**, The final subtomogram average map of P-RNAPII EC•RECQ5 complex from the condensates. RECQ5 IRI highlighted in yellow. **i**, Structural model of the in-condensate P-RNAPII EC•RECQ5 complex from subtomogram averaging (left) and its overlay with the cryo-EM off-condensate structure, RECQ5 IRI in violet (right).

We used cryo-ET as a single-molecule 3D imaging technique to determine the spatial organization of the P-RNAPII EC•RECQ5 condensates. We collected tomographic data and reconstructed 3D density maps of 105 condensates. We observed RNAPII condensates of different sizes with a spherical shape that was variable; some were rounded, whilst others were ellipsoidal, and some were at different stages of coalescence (Fig. 3c,d). We note that the largest RNAPII condensates were distorted in the blotting process, resulting in lentil-shaped condensates that were embedded within the thin ice layer. The median condensate volume was ∼0.005 μm^3^ (Fig. 3e), which matches that of the RECQ5 droplets formed at similar concentrations, as observed in the DLS experiments (Fig. 2e). It is also approximately five times smaller than the size of mediator condensates estimated by 3D super-resolution microscopy *in vivo*^45^. In addition to coalescence, the P-RNAPII EC particles were randomly oriented (Fig. 3f) and evenly distributed within the condensates (Fig. 3c), consistent with the liquid-like properties of the condensate.

Subtomogram averaging of particles within the condensates yielded a 7.1–Å–resolution cryo-ET map (Fig. 3h, Supplementary Table 1, Extended Data Fig. 6) into which we successfully fitted the cryo-EM structure of the RNAPII EC•RECQ5_1-650_ complex. We identified an additional density in the cryo-ET map that corresponds to a two-turn extension of the brake helix of RECQ5, which may enhance its contacts with the downstream DNA (Fig. 3i). Moreover, the cryo-ET map showed yet another, weak density near the RPB1 clamp and the RPB2 protrusion. Notably, the crystal structure of the RECQ5 helicase core^46^ matches the size of this additional density, and its position aligns with the anticipated size of the linker connecting the helicase to the brake helix (Extended Data Fig. 6d). We hypothesize that the helicase in this position might bind the single-stranded non-template strand before re-annealing it back to the template strand. The additional structural features seen in the P-RNAPII EC•RECQ5 complex inside the condensates could be attributed to the distinct chemical environment of the condensate, which may affect its structure and dynamics.

To fully reconstruct the P-RNAPII EC•RECQ5 condensate, elucidating the roles of the IDR and SRI domains of RECQ5, we used coarse-grained molecular dynamics (CG MD) simulations. We used the HPS-Urry parameterization, where each amino acid is represented by a single coarse-grained particle, to achieve the necessary timescales for condensate equilibration^47,48^. The HPS-Urry model was chosen due to its inclusion of phosphorylated serine. The model accurately reproduces experimentally determined properties of RECQ5 and P-RNAPII, such as ensemble-averaged radius of gyration and phase separation (Extended Data Fig. 7a,b). We added extra potentials to the rigid domains with the site-specific interactions (SRI with P-CTD and IRI with the RNAPII jaw) to match the experimental binding affinities (for details see SI, Extended Data Fig. 7e). In agreement with the experiments, our simulations showed condensation of CTD and RECQ5_IDR_ but no condensation of P-CTD (Extended Data Fig. 7b).

First, we investigated the self-association of RECQ5 in the absence of RNAPII, as a reference system (Extended Data Fig. 7d). In a simulation featuring 70 copies of RECQ5, a condensate spontaneously formed from a dispersed solution. On average, the RECQ5 IDR was found to be in contact with the IDR from 8±3 other RECQ5 copies (Extended Data Fig. 8c). An analysis of the inter-protein contacts (Fig. 4a) demonstrates the IDR’s significant contribution to molecular interactions. The cross-hatched pattern suggests that charged patches within the sequence drive these interactions.

**Fig. 4:**
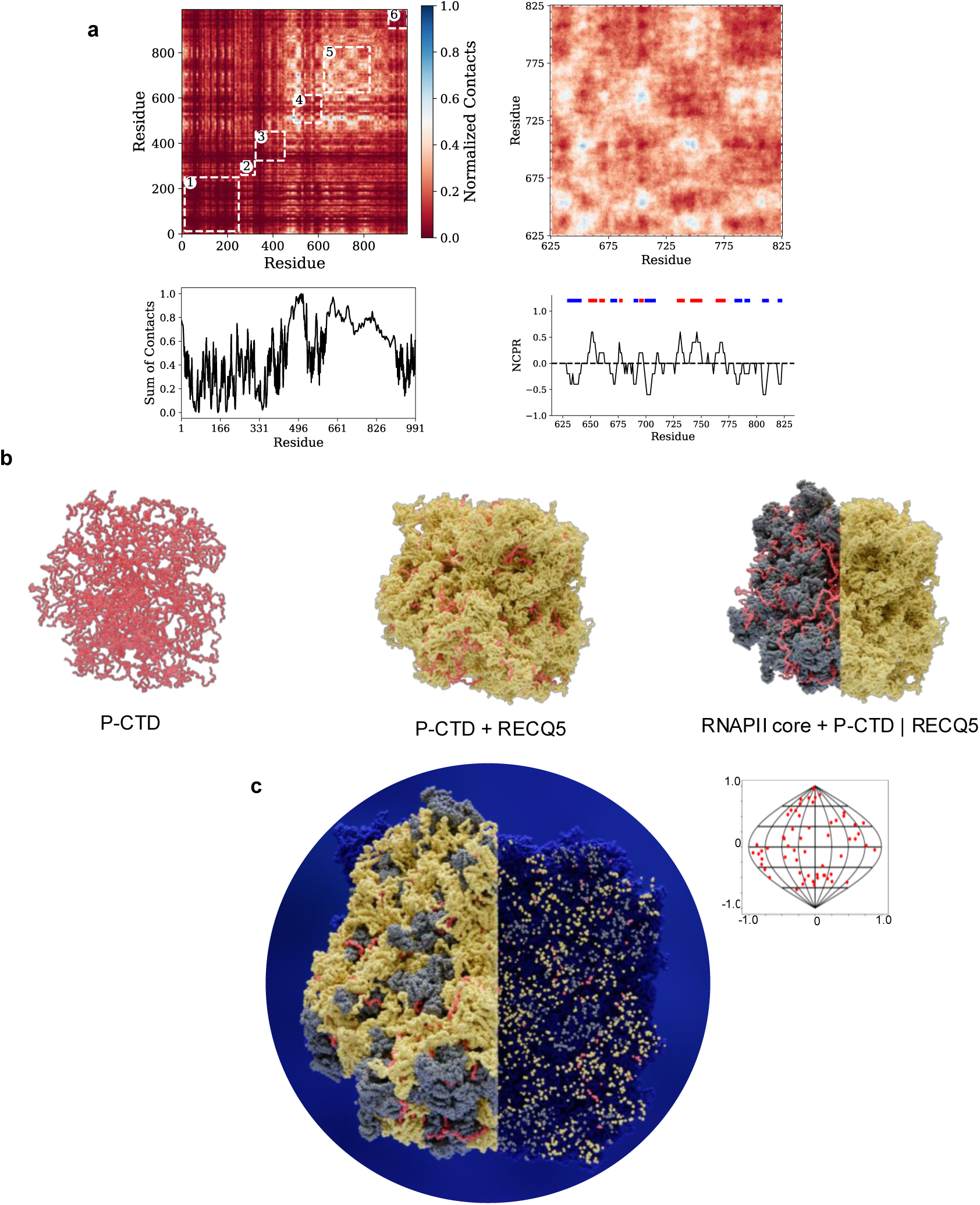
Molecular dynamics model of the P-RNAPII•RECQ5 condensate guided by experimental data. **a**, Inter-protein contact map of RECQ5 condensate. The heatmap is zoomed in for residues 625-825 (RECQ5_IDR_). The sum along one axis of the contact map reveals that the IDR is responsible for a large and continuous contact domain. This is likely due to the blocky charged region, which can be seen in the plot with net charge per residue (NCPR). **b**, Decomposition of the P-RNAPII•RECQ5 condensate. For an overview of the architecture, only P-CTD (in red, *left*), P-CTD and RECQ5 (in yellow, *middle*), P-CTD, RNAPII core (in grey) and RECQ5 (in a cross-sectional view, *right*) are shown. **c**, The complete view (*left* section) and a cross-sectional cut-through (*right* section) of the P-RNAPII•RECQ5 condensate, demonstrating occupied space by the proteins inside the condensate. Inlet, sinusoidal projection showing random P-RNAPII orientations in the simulated condensate.

To estimate the stoichiometry within the heterotypic P-RNAPII condensates, we simulated four small condensates of P-RNAPII and RECQ5 with P-RNAPII:RECQ5 ratios of 1:5, 1:7, 1:8, 1:9, respectively, using 10 copies of P-RNAPII. For all condensates, we calculated the separation distance between the closest P-RNAPII neighbors, in an attempt to best match the experimental results. Well-resolved P-RNAPII from cryo-ET maps of 70 condensates were used to calculate the minimum average separation distance of 12.6 nm. The same separation was best reproduced *in silico* in condensates with a ratio of 1:7 P-RNAPII:RECQ5 (Extended Data Fig. 8a). Subsequently, we carried out a simulation of a large condensate containing 60 copies of P-RNAPII and 420 RECQ5 molecules, to determine if the measured properties are size invariant.

The architecture of the equilibrated condensate revealed random distribution of the P-CTD of RNAPII and its core, as illustrated using decomposed views of the condensate (Fig. 4b). The orientations of P-RNAPII were randomly distributed, akin to our cryo-ET data (Fig. 4c). In both condensates containing 10 and 60 copies of P-RNAPII, respectively, between eight and nine copies of the RECQ5 SRI domain interacted with each P-CTD, on average (Extended Data Fig. 8d). Additionally, the average valency between RECQ5_IDR_ domains (Extended Data Fig. 8c) increased compared to the homotypic condensate which only contained RECQ5. Given current methods and computational limitations, determining the exact concentrations of RECQ5 and P-RNAPII using cryo-ET and MD simulations is unlikely. However, over all concentrations tested, interactions between RECQ5 and P-RNAPII consistently reinforced the structural stability of the condensate when compared to RECQ5 alone.

In summary, through the integration of cryo-ET imaging and MD simulations, we visualized the mesoscale organization of the RNAPII•RECQ5 molecular assembly and shed light on the underlying principles of its biogenesis.

## Discussion

RNAPII is thought to shuttle between types of transcription-associated condensates in a phosphorylation-dependent manner. Despite the continuing advancements in cellular cryo-ET, the molecular details of transcriptional condensates remain out of reach due to their small size and inherently dynamic nature. At present, our understanding of transcriptional condensates is based on superresolution imaging techniques^7,49,50^, which have inherent limitations to visualize the details in the spatial organization of these small condensates, as they offer only submicron resolved images. We used comprehensive biochemical analyses, single particle cryo-EM structure determinations, cryo-ET reconstitutions, *in vivo* validations, and coarse-grained simulations to study the molecular structure of the P-RNAPII-EC•RECQ5 condensate as a surrogate for transcriptional condensates existing *in vivo*. Through this integrative approach, we reconstructed the full structure of the P-RNAPII•RECQ5 condensate, uncovering critical interactions that shed light on the mechanisms of transcription attenuation and condensation.

Within the scaffold-client model of phase separation^51^, we show that RECQ5 undergoes phase separation driven by its IDR to form a scaffold matrix for P-RNAPII EC client molecules, which do not form condensates on its own when hyperphosphorylated. However, the recruitment of P-RNAPII EC into the RECQ5 scaffold matrix engages multivalent, site-specific interactions of the RECQ5 domains with the phosphorylated repetitive heptads of RNAPII CTD (52 heptads) and the downstream DNA in RNAPII’s jaws, thereby further reinforcing the integrity of the condensate (Fig. 5b). We propose that the phosphorylated repetitive heptad motif of the RNAPII CTD is crucial in enhancing the stability of condensates (Supplementary Fig. 6). It serves as a multivalent ligand platform and forms a complex network of site-specific interactions with scaffolding partners.

**Fig. 5.**
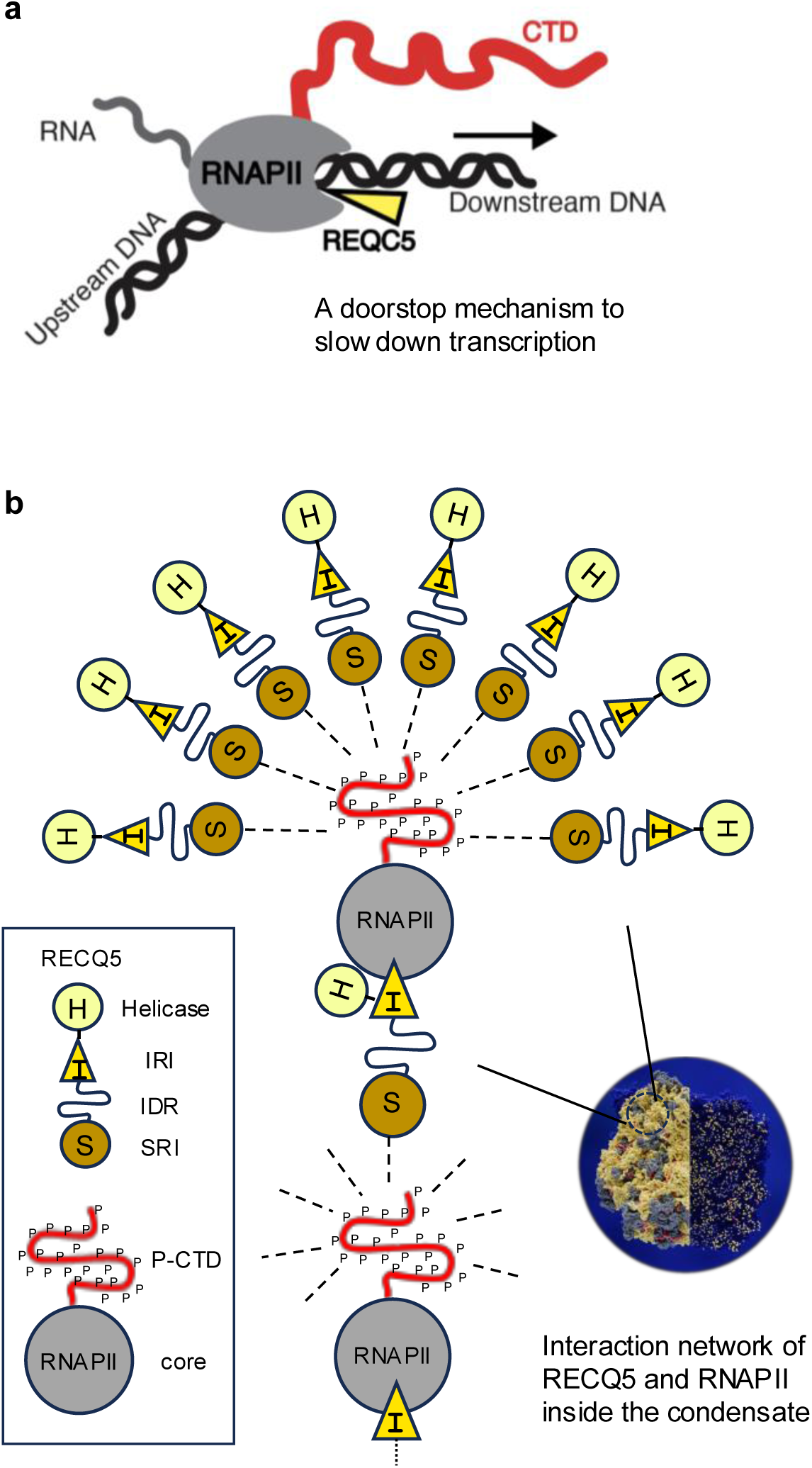
Model for transcription attenuation and condensation of RNAPII by RECQ5. **a**, Elongating RNAPII is slowed down by the IRI domain of RECQ5, which inserts its brake helix to contact downstream DNA entering RNAPII active center. **b**, Organization of RNAPII•RECQ5 condensates. RECQ5 undergoes phase separation driven by its IDR to form a scaffold matrix (on average one RECQ5 contacts 7 others) and integrates P-RNAPII client molecules via phosphorylation-dependent, site-specific interaction network, reinforcing the integrity of the P-RNAPII condensate.

Our cryo-EM structures and functional studies revealed the mechanism of how RECQ5 attenuates transcription of RNAPII. In contrast to all other structurally described regulatory factors, which associate with the upstream DNA during elongation^52–54^, RECQ5 associates with RNAPII’s jaw and inserts its brake helix to the jaw to directly contact the downstream DNA. Thus, the brake helix functions as a doorstop or physical barrier, modulating the movement of the RNAPII enzyme along the DNA (Fig. 5). The RNAPII binding sites for RECQ5 and TFIIS partially overlap, suggesting that their exchange is necessary to control elongation under different circumstances.

We speculate that not only physical contact of RECQ5 with downstream DNA entering the RNAPII catalytic center, but also the altered physicochemical properties in the condensate may be involved in slowing down transcript elongation rates via structural changes of RECQ5. Consistently, we showed that the structure of P-RNAPII EC•RECQ5 complex in condensate is changed at the brake helix, which is the critical component for the attenuation of transcription. Similarly, changes in the structural characteristics of other biomolecules, including nucleosomes, have been observed^55–57^. Our cryo-ET imaging showed that not only the hyperphosphorylated CTD, but also the entire RNAPII, including the transcribing core, are embedded within the condensates, addressing the key question of the spatial arrangement of transcriptional condensates. In summary, the results provide insight into RNAPII condensate assembly and lay the groundwork for future integrative studies of complex cellular condensates based on advanced cryo-ET 3D imaging combined with coarse-grained simulations.

## Methods

### Plasmids

Plasmids pTXB1-RECQ5, pTXB1-RECQ5^K58R^ were generously provided by Pavel Janscak (University of Zurich)^58^. Plasmids pTXB ^K58R, L556E, L602E,^ pTXB1-RECQ5^K58R, R943A^, pTXB1-RECQ5_1-625_, pTXB1-RECQ5_1-650_, pTXB1-RECQ5_1-650_^K58R^, pTXB1-RECQ5_1-650_^K58R, L556E, L602E,^ pTXB1-RECQ5_1-650_^K58R, R502A,^ pTXB1-RECQ5_1-650_^K58R, R502E,^ and pTXB1-RECQ5_1-650_^K58R, Δ490–515(BH),^ were generated by PCR. To express and purify RECQ5 and its fragments (RECQ5_625-901_, RECQ5_625-825_) fused with a fluorescent tag, fragment of DNA containing the ORF was cloned into plasmids H6-mCerulean (pET Biotin His6 TEV mCerulean LIC cloning vector, Addgene plasmid #29726), yielding in-frame fusion of RECQ5 with mCerulean. Next, the fused ORFs were subcloned into vector 438B (pFastBac His6 TEV cloning vector with BioBrick Polypromoter LIC subcloning, Addgene plasmid #55219), enabling transposition into bacmids (438B-RECQ5-mCerulean). Plasmids 438B-RECQ5^ΔSRI^-mCerulean, 438B-RECQ5^Δ515-620(KIX)-^mCerulean, 438B-RECQ5^Δ625-901^-mCerulean, 438B-RECQ5^Δ625-825^-mCerulean, 438B-RECQ5^Δ675-775^-mCerulean, and 438B-RECQ5^Δ625-825ΔSRI^-mCerulean were generated by PCR. To express and purify fragments RECQ5_625-901_, RECQ5_625-825_, and RECQ5_475-901_, fragments of DNA containing the coding sequence were amplified by PCR and cloned into vector 2BT using LIC. To express RECQ5 fused with mGFP in human cell-lines, the ORF was cloned into plasmid 6D (pcDNA3 GFP LIC cloning vector, Addgene plasmid **#**30127). Mutant versions 6D-RECQ5^ΔSRI^, 6D-RECQ5^ΔKIX^, 6D-RECQ5^Δ625-825^, and 6D-RECQ5^Δ625-825ΔSRI^ were generated by PCR.

To generate plasmids enabling expression of the kinase module of TFIIH complex (the CDK7 complex) in insect cells, the ORFs for CDK7, MAT1, and CCNH were individually cloned into plasmid 438B. Plasmids 438B-CDK7, 438B-MAT1, and 438B-CCNH were subsequently combined using BioBrick Polypromoter LIC subcloning^59^ into a single construct, enabling co-expression of the three subunits of the complex from a single virus in insect cells.

To express and purify the full-length, C-terminal domain of the catalytic subunit of RNAPII (hCTD) fused with mCherry, fragment of DNA containing codon-optimized ORF of mCherry-hCTD was cloned into vector 2BcT (pET His6 LIC cloning vector, Addgene plasmid #37236). To express and purify the JAW domain of RPB1 (AA 1168-1302), the fragment of DNA, containing the coding sequence was amplified by PCR and cloned into 2GT vector by LIC.

Plasmid pGEX4T1-(CTD)_26_-(His)_7_ was used to express and purify GST-(CTD)_26_-(His)_7_. Plasmids 2BT, 2BcT, 2GT, 6D, 438B, H6-mCerulean, and H6-mCherry were purchased directly from QB3 Macrolab (UC Berkeley). All constructs were verified by sequencing.

A list of oligonucleotides is listed in Supplementary Table 2, detailed information on the vectors used is listed in Supplementary Table 3, and a list of prepared recombinant plasmids is provided in Supplementary Table 4.

### Cell culture

The HEK293T cells (ATCC-CRL-11268) were grown at 37 °C and 5% CO_2_ in complete DMEM medium (Gibco #41966-029), supplemented with 10% fetal bovine serum, 2 mM L-glutamine (Life Technologies, #25030024), 50 U/ml penicillin, and 50 U/ml streptomycin (Hyclone-Biotech, #SV30010). Cell passaging was performed using trypsin (Biosera). Cells were washed in Phosphate Buffered Saline (PBS) buffer prior to the passage.

### Transient Transfection

Adherent cells were seeded at a density of 10000 cells per cm^2^ and were incubated for 24 h. The cells were transfected using Polyetylenimin (PEI) (1 mg/ml stock concentration, adjusted to pH 7.4). The amount of plasmid DNA used for transfection was 300 ng/cm^2^, and the ratio of PEI and DNA was 4:1. Firstly, DNA and PEI were separately mixed with serum-free DMEM, incubated for 20 min at room temperature (RT) and subsequently mixed. Each mixing step was followed by vigorous shaking and centrifugation. The final mix was incubated for 20 min at RT and added to the cells. After 4-hour incubation with the transfection mixture, the medium was replaced with fresh complete DMEM. The cells were either analyzed or fixed 24 hours after transfection.

### Immunofluorescence and live-cell imaging

For live cell imaging, HEK293T cells were seeded onto µ-Slide 8 Well (Ibidi) chambers. For fixed samples HEK293T cells were seeded onto 13mm coverslips in 24-well plate, coated in 0.1% gelatine. The next day, the cells were transfected with plasmids as described above and subsequently incubated for 24 hours. Then live cell imaging was performed using confocal (Leica SP8) microscope equipped with culture chamber at 37 °C and 5% CO_2_. Micrographs were taken using oil immersion objective with 60x magnification. For fixed samples, cells were washed in PBS, fixed in 4% paraformaldehyde (PFA, Millipore) in PBS for 15 min, followed by three washes in PBS. Subsequently, the cells were permeabilized by 0.3% Triton X-100 in PBS for 15 min, and finally blocked in PBTA (PBS, 5% donkey serum, 0.3% Triton X-100, 1% BSA) for 1 hour. Samples were incubated overnight at 4°C with primary antibody (pSer5-CTD mAb (rat, clone#3E8)) diluted in PBTA in ratio 1:200. Following 3 washes in PBS, the samples were incubated with the corresponding Alexa Fluor 568 secondary antibody with dilution in PBTA 1:500 (Invitrogen, # A-11077) for 2 hours at RT. Then, the cells were washed 3x in PBS, followed by 10 min incubation with DAPI (Thermo Fisher) at RT for nuclear staining. Finally, samples were washed once in PBS and mounted in DAKO KO mounting solution (DAKO KO). The micrographs were taken using confocal (Leica SP8) microscope using oil immersion objective with 60x magnification.

#### Data analyses

For protein subcellular localization experiments, at least 100 positive cells per experiment (n = 3) were analyzed and scored according to their phenotype into two categories (condensates/even).

### Insect cell work

To generate viruses enabling production of proteins in insect cells, the coding sequences and the necessary regulatory sequences of the constructs were transposed into bacmid using *E. coli* strain DH10bac (Thermofisher). The viral particles were obtained by transfection of the bacmids into the *Sf*9 cells using FuGENE Transfection Reagent (Promega) and further amplification in *Sf*9 cells. Proteins were expressed in 200 ml of Hi5 cells (infected at 1×10^6^ cells/ml) with the corresponding P1 virus at multiplicity of infection >1. The cells were harvested 48 hours post infection, washed with 1x PBS, and stored at −80°C.

### Protein purification

#### Purification of RECQ5 and its variants

RECQ5 and its mutant versions were purified as described^60^ with modifications. Briefly, RECQ5 was purified from cleared cell-lysate by successive affinity purifications over chitin-beads, heparin column, followed by size-exclusion chromatography on Superdex S-200 column equilibrated with 25 mM Tris-Cl, pH7.5, 300 mM NaCl, 1 mM DTT. Fractions containing RECQ5 were pooled, concentrated, snap-frozen in liquid nitrogen, and stored at −80 °C.

#### Purification of RECQ5_625-901_, RECQ5_625-825_, RECQ5_475-901_, RECQ5_625-901_-mCerulean, RECQ5_625-825_-mCerulean, and mCerulean

Ten grams of *E. coli* BL21 RIPL cells expressing RECQ5_625-901_, RECQ5_625-825_, RECQ5_475-901_, RECQ5_625-901_-mCerulean, RECQ5_625-825_-mCerulean, and mCerulean, respectively, were resuspended in ice-cold lysis buffer (50 mM Tris-HCl, pH 8, 0.5 M NaCl, 10 mM imidazole, 1 mM DTT) containing protease inhibitors (0.66 μg/ml pepstatin, 5 μg/ml benzamidine, 4.75 μg/ml leupeptin, 2 μg/ml aprotinin) at +4°C. Cells were opened up by sonication. The cleared lysate was passed through 2 ml of Ni-NTA beads (Qiagen), equilibrated with 50 mM Tris-HCl, pH 8, 500 mM NaCl, 10 mM imidazole, and 1 mM DTT. The proteins were eluted with an elution buffer containing 50 mM Tris-HCl, pH 8, 500 mM NaCl, 1 mM DTT, and 400 mM imidazole. The elution fractions containing fragments of RECQ5 were pooled and dialyzed for 3 hours against 50 mM Tris-HCl, pH 8, 500 mM NaCl, 10 mM imidazole, and 1 mM DTT in the presence of TEV protease to cleave the His_6_-tag off. Subsequently, the dialysate was passed through 1 ml of Ni-NTA beads, concentrated, and further fractioned on Superdex-S75 column with SEC buffer (25 mM Tris-Cl, pH7.5, 200 mM NaCl, 1 mM DTT). Fractions containing homogeneous proteins were concentrated, snap-frozen in liquid nitrogen, and stored at −80 °C.

#### Purification of RECQ5-mCerulean and its variants

Pellets of Hi5 insect cells, expressing RECQ5-mCerulean and its mutant variants were resuspended in ice-cold lysis buffer containing 50 mM Tris-Cl, pH 8.0, 500 mM NaCl, 0.4% Triton X-100, 10% (v/v) glycerol, 10 mM imidazole, 1 mM DTT, protease inhibitors (0.66 μg/ml pepstatin, 5 μg/ml benzamidine, 4.75 μg/ml leupeptin, 2 μg/ml aprotinin), and 25 U benzonase per ml of lysate. The resuspended cells were gently shaken for 10 min at 4°C. To aid the lysis, cells were briefly sonicated. The cleared lysate was passed through 2 mL of Ni-NTA beads (Qiagen), equilibrated with 50 mM Tris-Cl, pH 8, 500 mM NaCl, 10 mM imidazole, and 1 mM DTT. Proteins were eluted with an elution buffer containing 50 mM Tris-Cl, pH 8, 500 mM NaCl, 1 mM DTT, and 400 mM imidazole. The elution fractions containing proteins were pooled and dialyzed overnight against 50 mM Tris-Cl, pH 8, 500 mM NaCl, 10 mM imidazole, and 1 mM DTT in the presence of TEV protease to cleave the His_6_-tag off. The next day, the dialysate was passed through 1mL of Ni-NTA beads, concentrated, and further fractioned on Superose 6 column with SEC buffer (25 mM Tris-Cl pH 7.5, 200 mM NaCl, 1 mM DTT). Fractions containing homogeneous proteins were concentrated, snap-frozen in liquid nitrogen, and stored at −80 °C.

#### Purification of the CDK7 complex

Pellets of Hi5 insect cells, expressing the CDK7 complex were resuspended in ice-cold lysis buffer containing 50 mM Tris-Cl pH 8.0, 500 mM NaCl, 0.4% Triton X-100, 10% (v/v) glycerol, 10 mM imidazole, 1 mM DTT, protease inhibitors (0.66 μg/ml pepstatin, 5 μg/ml benzamidine, 4.75 μg/ml leupeptin, 2 μg/ml aprotinin), and 25 U benzonase per ml of lysate. The resuspended cells were gently shaken for 10 min at 4°C. To aid the lysis, the cells were briefly sonicated. The cleared lysate was passed through 2 ml of Ni-NTA beads (Qiagen), equilibrated with 50 mM Tris-Cl, pH 8, 500 mM NaCl, 10 mM imidazole, and 1 mM DTT. Proteins were eluted with an elution buffer containing 50 mM Tris-Cl, pH 8, 500 mM NaCl, 1 mM DTT, and 400 mM imidazole. The elution fractions containing proteins were pooled, concentrated, and further fractioned on Superdex S-200 column with SEC buffer (25 mM Tris-Cl pH7.5, 200 mM NaCl, 1 mM DTT). Fractions containing homogenous complex were concentrated, glycerol was added to a final concentration of 10 % before they were snap-frozen in liquid nitrogen, and stored at −80 °C.

#### Purification of mCherry-hCTD

Ten grams of *E. coli* BL21 RIPL cells expressing mCherry-hCTD were resuspended in ice-cold lysis buffer (50 mM Tris-Cl, pH 8, 500 mM NaCl, 10 mM imidazole, 1 mM DTT) containing protease inhibitors (0.66 μg/ml pepstatin, 5 μg/ml benzamidine, 4.75 μg/ml leupeptin, 2 μg/ml aprotinin) at +4°C. Cells were opened up by mild sonication, followed up by cell cracking. The cleared lysate was passed through 2 ml of Ni-NTA beads (Qiagen), equilibrated with 50 mM Tris-Cl, pH 8, 500 mM NaCl, 10 mM imidazole, and 1 mM DTT. mCherry-hCTD was eluted with an elution buffer containing 50 mM Tris-Cl, pH 8, 500 mM NaCl, 1 mM DTT, and 400 mM imidazole. The elution fractions containing mCherry-hCTD were pooled and dialyzed overnight against 50 mM Tris-Cl, pH 8, 500 mM NaCl, 10 mM imidazole, and 1 mM DTT in the presence of TEV protease to cleave the His_6_-tag off. The next day, the dialysate was passed through 1ml of Ni-NTA beads, concentrated, and further fractioned on Superdex-S200 column with SEC buffer (25 mM HEPES-OH pH7.5, 220 mM NaCl, 1 mM DTT). Fractions containing homogeneous proteins were concentrated, snap-frozen in liquid nitrogen, and stored at −80 °C.

#### Purification of mammalian RNAPII

The mammalian, twelve-subunit RNAPII complex was purified from calf (*Bos taurus*) thymus. Briefly, chromatin-bound RNAPII was purified to homogeneity by successive rounds of ion-exchange, immuno-affinity, ion-exchange, and size-exclusion chromatography as described in^61^.

#### Purification of phosphorylated mammalian RNAPII

To generate RNAPII phosphorylated on serine 5 and 7 (P-RNAPII), the procedure was performed as described for RNAPII, with modifications. The pooled sample from the second ion-exchange chromatography (MonoQ), was concentrated to ∼500µL. To the concentrated sample, the CDK7 complex (100 µg) and a mixture of ATP/MgCl_2_ (at a final concentration 2 mM and 3.5 mM, respectively) was added. The volume was adjusted to 1 mL, and the mixture was incubated at 30°C for 30 min. Subsequently, the P-RNAPII was separated from the reagents on Superose 6 size-exclusion column. Fractions containing P-RNAPII were pooled, concentrated, snap-frozen in liquid nitrogen, and stored at −80°C.

#### Purification of GST-JAW(AA1168-1302) domain of RPB1

Ten grams of *E. coli* BL21 RIPL cells expressing GST-JAW were resuspended in ice-cold lysis buffer (50 mM Tris-Cl, pH 8, 500 mM NaCl, 10 mM imidazole, 1 mM DTT) containing protease inhibitors (0.66 μg/ml pepstatin, 5 μg/ml benzamidine, 4.75 μg/ml leupeptin, 2 μg/ml aprotinin) at +4°C. Cells were opened up by sonication. The cleared lysate was passed through 2 mL of Ni-NTA beads (Qiagen), equilibrated with 50 mM Tris-Cl, pH 8, 500 mM NaCl, 10 mM imidazole, and 1 mM DTT. GST-JAW was eluted with an elution buffer containing 50 mM Tris-Cl, pH 8, 500 mM NaCl, 1 mM DTT, and 400 mM imidazole. The elution fractions containing GST-JAW were pooled and applied onto Superdex-S200 column with SEC buffer (25 mM Tris-Cl pH7.5, 200 mM NaCl, 1 mM DTT). Fractions containing homogeneous protein were concentrated, snap-frozen in liquid nitrogen, and stored at −80 °C.

#### Preparative phosphorylation and purification of CTD polypeptides

For preparative purposes, 2.5 mg of GST-(CTD)_26_-(His)_7_ and mCherry-hCTD, respectively, were phosphorylated by 250 µg of the CDK7 complex (to phosphorylate serine 5 and serine 7 on the CTD) or by 250µg of c-Abl (to phosphorylate tyrosine 1 on the CTD) in the presence of 2 mM ATP and 3.5 mM MgCl_2_ for 60 min at 30°C. Reactions were stopped by placing at +4°C. The CTD peptides were purified from the kinases and ATP by size-exclusion chromatography on Superdex S-200, equilibrated with 25 mM Tris-Cl, pH 7.5, 220 mM NaCl, 1 mM DTT. Fractions containing phosphorylated CTD polypeptides were pooled, concentrated, snap-frozen in liquid nitrogen, and stored at −80°C.

### *In vitro* LLPS assays

Liquid-liquid phase separation (LLPS) assays were performed in buffer H_220_ (25 mM HEPES-OH, pH 7.5, 220 mM NaCl, 0.5 mM TCEP), where indicated, crowding agent was present (3.75% (w/v) PEG-8000, unless stated otherwise). Where ssDNA was used, it was added at 1:10 ratio (Cy3-labeled: unlabeled) in the absence of the crowding agent in buffer H_150_ (25 mM HEPES-OH, pH 7.5, 150 mM NaCl, 0.5 mM TCEP). Upon addition of the indicated proteins (RECQ5-mCerulean variants at 1.25, 2.5, and 5 µM, except for RECQ5_625-901_-mCerulean, RECQ5_625-825_-mCerulean, which were present at 2.5, 5, and 10 µM, mCherry-P-CTD at 1.25, 2.5, and 5µM, mCerulean at 2.5, 5, 10, and 20 µM), the mixtures were immediately spotted onto a glass slide, and the condensates were recorded on Zeiss Axio Observer.Z1 with a 63x water immersion objective.

In the experiments in which the elongation complex (EC) of P-RNAPII was used, the ECs were first assembled by mixing 2pmols of either unmodified RNAPII or P-RNAPII with 1.8pmols of the annealed RNA:DNA hybrid, containing the template strand and Cy-3 labelled RNA, followed by incubation for 20 min at 30°C. Subsequently, the non-template strand (2pmols) was added, and the mixture was incubated for further 20 min at 30°C. The assembled ECs of (P-)RNAPII (at 250 nM in 10 mM HEPES-OH pH7.5, 100 mM NaCl, 10 mM DTT) were mixed with 500 nM RECQ5-mCerulean, RECQ5^ΔKIX^-mCerulean, RECQ5^ΔSRI^-mCerulean, and RECQ5^ΔIDR^-mCerulean, respectively. Prior to visualization, PEG8000 was added to a final concentration 3.75% (w/v). The samples were subsequently visualized as described above.

#### Statistical analyses

Analyses and quantifications of the micrographs were performed in Cell-Profiler (version 4.2.1)^62^. First, five micrographs (2048 pixels (px) per 2048 px, 1 px= 0.103174 µm) per condition and per experiment were analyzed. Objects (droplets) were identified based on diameter (2-70 px, 0,206-7.5 µm) and intensity using Otsu’s method for thresholding^63^. Picked objects were further filtered based on eccentricity and intensity. For the filtered objects the area, median intensity of the objects for each channel (mCerulean, mCherry/Cy3), and the object count per pictures were calculated. R^64^ and R-studio^65^ with tidyverse package^66^ were used to process the data obtained from Cell-profiler analysis. The values for droplets areas were converted from the px to µm based on the metadata of the micrographs (0.010645 µm^2^ = 1px). Graphs for the figures were plotted using ggplot2^67^, ggbeeswarm^68^, ungeviz^69^, and ggpubr^70^ packages. Statistical analysis for the area size comparison was done using unpaired t-test (*14*). ns p>0.05, *p≤ 0.05, **p≤ 0.01, ***p≤ 0.001, ****p≤ 0.0001.

### *In vitro* transcription assay

The dC tailed *H3F3A* (encoding histone H3.3) intron template was prepared by digestion of the pGEMTerm plasmid (kindly provided by Susanne Kassube) as described previously^71–73^. Reactions (15 μl) in transcription buffer (25 mM HEPES-OH pH 7.9, 100 mM KCl, 5 mM MgCl_2_, 5% glycerol, 6 mM spermidine, 1 mM DTT, 1U RNasin Plus, 800 μM each of GTP, UTP, and ATP) contained 20 μg/ml dC-pGEMTerm, 0.8 pmol mammalian RNAPII, 1 or 10 pmol of RECQ5 variants (as indicated in the figures), or the equivalent amount of buffer. After the addition of 1 μl [α-^32^P]-CTP (0.01 μM), the reactions were incubated for 3 min at 30°C. Following the addition of 3.5 μl of the chase buffer (25 mM HEPES-OH pH 7.9, 100 mM KCl, 5 mM MgCl_2_, 1 mM DTT, 800 μM each of GTP, UTP, ATP, and 4.8 mM CTP), incubation at 30 °C was continued. After 10 min, the samples were digested with proteinase K for 10 min at 30°C before the addition of formamide loading buffer (0.02% SDS, bromophenol blue in 90% formamide). RNA transcripts were resolved on a 6% polyacrylamide gel containing 7 M urea and visualized by phosphorimaging.

### *In vitro* pull-down experiments

Purified GST-CTD, GST-Y1P-CTD, or GST-S5,7P-CTD (5 μg each), respectively, were incubated with RECQ5 (5 μg) in 30 μl of buffer T_200_ (25 mM Tris–Cl pH7.5, 200 mM NaCl, 10 % glycerol, 1 mM DTT, 0.5 mM EDTA, and 0.01% Nonidet P-40) for 30 min at 4°C in the presence of GSH-beads. After washing the beads twice with 100 μl of buffer T_200_, the bound proteins were eluted with 30 μl of 4xSDS loading dye. The input, supernatant, and eluate, 7 μl each, were analyzed on SDS-PAGE gel. Alternatively, purified GST-JAW (5 µg) was incubated with RECQ5_1-625_, RECQ5_1-650_ (and its mutant variants) (5 μg each) in 30 µL of buffer T_150_ (20 mM Tris–Cl pH 7.5, 150 mM NaCl, 10 % glycerol, 1 mM DTT, 0.5 mM EDTA, and 0.01% Nonidet P-40) for 30 min at 4°C in the presence of GSH-beads. Subsequently, the reactions were processed as described for pull-down experiments with GST-CTDs.

### Electrophoretic Mobility-Shift Assay (EMSA)

Increasing concentrations of the tested proteins (at 12.5, 25, 50, and 100 nM, respectively) were incubated with fluorescently labelled ssDNA (61-mer, final concentration 10 nM) in buffer D (25 mM Tris-Cl, pH 7.5, 1 mM DTT, 5 mM MgCl_2_, and 100 mM NaCl) for 20 min at 37°C. Loading buffer (60 % glycerol in 0.001% Orange-G) was added to the reaction mixtures and the samples were loaded onto a 7.5 % (w/v) polyacrylamide native gel in 0.5 x TBE buffer and run at 75 V for 1h at +4°C. The different nucleic acid species were visualized using FLA-9000 Starion scanner and quantified in the MultiGauge software (Fujifilm). To calculate the relative amount of bound nucleic acid substrate, the background signal from the control sample (without protein) was subtracted using the *band intensity - background* option. Nucleic acid-binding affinity graphs were generated with Prism-GraphPad 7.

### *In vitro* crosslinking experiments

To determine the oligomeric state of RECQ5_625-901_ and RECQ5_625-825_, the proteins were diluted to 0.5 mg/ml in buffer H_150_ (25 mM HEPES pH7.5, 150 mM NaCl, 1 mM DTT). To the diluted proteins, glutaraldehyde (0.05% final concentration) was added, and the mixture was incubated for 30 min at 4°C. Subsequently, the reactions were stopped by the addition of 1µl of 1M Tris-Cl. The reactions were analyzed by SDS-PAGE.

### Dynamic light scattering (DLS)

The DLS measurements were carried out on a Delsa Max Core (Beckman Coulter) instrument. RECQ5 (0.1, 0.15, 0.2, 0.3, and 0.4 µM) in the buffer H_150_ was mixed with 2µM Cy3-labeled ssDNA to promote liquid-liquid phase-separation. The resulting reaction mixture (10 µL) was transferred to a disposable Wyatt-type cuvette and the measurement was started 30s after the components had been mixed. The intensity of the scattered light was recorded at a 90° scattering angle, at 25°C, and a wavelength of 662 nm. Each individual time-course experiment consisted of 240 acquisitions (5 seconds each).

The software package DelsaMax Analysis Software 1.0 was used to visualize and process the acquired data. Hydrodynamic radii were calculated through cumulant fitting of the autocorrelation function and were plotted against time. Every measurement for all RECQ5 concentrations was done in triplicates. R and R-studio with tidyverse package were used to plot the data obtained from DelsaMax Analysis Software 1.0. Graphs for the figures were plotted using ggplot2. The plotted dots represent the average values from three measurements, with error bars indicating the standard deviation. Highlighted is every 10^th^ point from the measurements. Data were fitted with y=log(x) function.

### Small-angle X-ray scattering (SAXS)

SAXS data from solutions of wild type RECQ5 and RECQ5_IDR_ were collected using a Rigaku BioSAXS-2000 instrument at CEITEC (Brno, Czech Republic) equipped with a HyPix-3000 detector at a sample-detector distance of 0.5 m (I(s) vs s, where = 4π sin θ/λ; 2θ is the scattering angle and λ = 0.154 nm). Six successive 3600 second frames were collected at a sample temperature of 20°C using the following concentration series: RECQ5: 1, 2, and 4 mg/ml; RECQ5_IDR_: 0.2, 0.4, 0.6, 0.8, and 1 mg/ml. The data were normalized to the intensity of the transmitted beam and radially averaged and the corresponding scattering from the solvent-blank was subtracted to produce the scattering profile displayed. The radii of gyration (R*_g_*) were determined by Autorg as is implemented in Primusqt^74^.

### Correlative light-electron microscopy (CLEM)

#### Sample preparation

The elongation complex (EC) of P-RNAPII was first assembled by mixing 2 pmols of P-RNAPII with 2.5 pmols of the annealed RNA:DNA hybrid, containing the template strand and FITC-labelled RNA, followed by incubation for 20 min at 30°C. Subsequently, the non-template strand (3 pmols) was added, and the mixture was incubated for further 20 min at 30°C. The sample of 100 nM EC of P-RNAPII were preincubated with 250 nM RECQ5-mCerulean in RNAPII buffer (10 mM HEPES-OH pH 7.5, 100 mM NaCl, 10 µM ZnCl_2_, 2mM MgCl_2_, and 2 mM DTT) for 15 min. Subsequently, glutaraldehyde (0.1% final concentration) was added, and the mixture was incubated on ice for further 15 minutes.

#### Data acquisition

Sample of the EC of P-RNAPII•RECQ5-mCerulean complex (3.5 µL) was applied to freshly glow-discharged Au 300 R1.2/1.3 TEM grid (Quantifoil) coated with a monolayer of graphene and vitrified into liquid ethane using Vitrobot Mark IV (ThermoScientific) (5°C, 100% rel. humidity, 30 s waiting time, 4 s blotting time). The grids were subsequently mounted into the Autogrid cartridges and loaded to Talos Arctica (ThermoScientific) transmission electron microscope operated at 200 kV.

A low magnification (155x) hole grid montage was collected in SerialEM, followed by acquisition of the selected grid squares at 900x magnification. The grid was subsequently transferred into DMI6 cryo-CLEM (Leica) wide-field microscope and hole grid z-stack montages of the grids were collected with a 50x objective under the cryo-conditions in transmission light, DAPI (for mCerulean), and GFP (for FITC-labeled EC of P-RNAPII) channels. The data from the wide-field microscope was processed and analyzed using LAS X CLEM software (Leica) and the two types of the data were manually overlayed in Fiji.

### Cryo-electron microscopy

#### Sample preparation

The elongation complexes (ECs) were first assembled by mixing 2 pmols of either unmodified RNAPII or P-RNAPII with 2.5 pmols of the annealed RNA:DNA hybrid, containing the template strand and FITC labelled RNA, followed by incubation for 20 min at 30°C. Subsequently, the non-template strand (3 pmols) was added, and the mixture was incubated for further 20 min at 30°C. In the experiments containing P-RNAPII EC and RECQ5^K58R^, P-RNAPII EC and RECQ5^ΔIDR^, RNAPII EC and RECQ5_1-650_, RNAPII EC and RECQ5_1-650_^R502E^ a mixture of 2 mM CTP and 3.5 mM MgCl_2_ was added alongside the non-template strand.

Sample of 100 nM (EC) of P-RNAPII was preincubated with 250 nM RECQ5^K58R^ for 15 min in RNAPII buffer. Subsequently, glutaraldehyde (0.1% final concentration) was added, and the mixture was incubated on ice for a further 15 minutes. Alternatively, 1 µM EC of RNAPII was preincubated with 5 µM RECQ5_1-650_, RECQ5_1-650_^R502E^, RECQ5^ΔIDR^, and RECQ5^K58R^, respectively, for 15 min in RNAPII buffer. Subsequently, the mixture was diluted 10-fold and glutaraldehyde (0.1% final concentration) was added, followed by incubation for further 15 min on ice.

The mixture (3.5 µL) was applied to freshly glow-discharged Au 300 R1.2/1.3 TEM grids (Quantifoil) coated with the monolayer of graphene and vitrified into liquid ethane using Vitrobot Mark IV (ThermoScientific) (5°C, 100% rel. humidity, 30 s waiting time, 6 s blotting time). The grids were subsequently mounted into the Autogrid cartridges and loaded to Talos Arctica or Titan Krios (ThermoScientific) transmission electron microscopes (TEMs) for the data acquisition.

#### Data acquisition

##### RNAPII EC•RECQ5^K58R^ complex (off-condensate particles)

The RNAPII EC•RECQ5^K58R^ complex data were collected using Talos Arctica (TEM) operated at 200 kV using SerialEM software^75^. The data were acquired using Falcon3 (ThermoScientific) direct electron detector operated in linear mode. Micrographs were collected at the calibrated pixel size of 1.22 Å/px as a set of 40 frames comprising the overall dose of 40 e^-^/Å^2^. The complete dataset consisted of 2552 movies.

##### RNAPII EC•RECQ5^ΔIDR^ complex

The RNAPII EC•RECQ5^ΔIDR^ data were collected using Talos Arctica (TEM) as described above. The energy selecting slit was set to 15 e^-^V. Micrographs were collected at the calibrated pixel size of 0.783 Å/px as a set of 30 frames comprising the overall dose of 45 e^-^/Å^2^. The complete dataset consisted of 5724 movies.

##### RNAPII EC•RECQ5_1-650_ complex

The RNAPII EC•RECQ5_1-650_ complex data were collected using Titan Krios (TEM) operated at 300 kV using SerialEM software^75^. The data were acquired using Bioquantum K3 (Gatan) direct electron detector operating in electron counting mode and positioned behind the Gatan Imaging Filter. The energy selecting slit was set to 10 e^-^V. Micrographs were collected at the calibrated pixel size of 0.8336 Å/px as a set of 40 frames comprising the overall dose of 40.2 e^-^/ Å^2^. The complete dataset consisted of 12410 movies.

##### RNAPII EC•RECQ5_1-650_ ^R502E^ complex

The RNAPIIEC•RECQ5_1-650_ ^R502E^ data were collected using Titan Krios (TEM) as described above. The energy selecting slit was set to 10 e^-^V. Micrographs were collected at the calibrated pixel size of 0.8336 Å/px as a set of 40 frames comprising the overall dose of 36.6 e^-^/Å^2^. The complete dataset consisted of 9046 movies.

#### Data processing

##### RNAPII EC•RECQ5^K58R^ complex (off-condensate particles)

The movies were imported into cryoSPARC 3.2^76^ and aligned using the patch motion correction job. The CTF parameters were estimated with Patch CTF job. The images with the estimated CTF fit worse than 7 Å were excluded from further data analysis. A set of 10 micrographs was randomly selected from the dataset for manual particle-picking to generate references for the template picking of the whole dataset. The overall number of 678,625 particles was extracted from the micrographs and the false positive picks and corrupted particles were removed by running multiple rounds of reference-free 2D classification resulting in 189,964 particles, which were used to for the downstream data analysis. The *ab-initio* job was used to generate two initial models and carry out the first 3D classification of the data. Models corresponding to RNAPII EC in apo from and bound to RECQ5 were obtained at 3.6 and 3.8 Å, respectively (according to the FSC = 0.143 criterion).

##### RNAPII EC•RECQ5^ΔIDR^ complex

The movies acquired with Talos Arctica (TEM) were imported into cryoSPARC 4.02^76^ and aligned using the patch motion correction job. The CTF parameters were estimated with Patch CTF job. The images with the estimated CTF fit worse than 7Å were excluded from further data analysis. A set of 20 micrographs was randomly selected from dataset for manual particle-picking to generate references for the template picking of the whole dataset. The overall number of 223,895 particles was extracted from the micrographs and binned 2x to 128 px box size (2.40 Å/px). The false positive picks and corrupted particles were removed by reference-free 2D classification resulting in 181,882 particles, which were used for the downstream data analysis. The *ab-initio* job was used for the initial 3D classification of the data. A model corresponding to RNAPII•RECQ5^ΔIDR^ complex, which contained 147,368 particles was further refined and those particles were re-extracted to 352 px boxsize (0.783 Å/px). Another *ab-initio* job was used to generate initial model. *NU-Refinement* followed by the CTF refinement resulted in a 3.2 Å RNAPII•RECQ5^ΔIDR^ map. The cryo-EM density of RECQ5 and the active site was used to create a local mask. A classification strategy involving initial classification of RECQ5, followed by focused classification of the active site, was employed to identify sub-populations of 15168 particles with well-resolved densities for both the active site and RECQ5. The final map of RNAPII•RECQ5^ΔIDR^ had resolution of 3.8 Å (according to the FSC = 0.143 criterion). The refinement statistics are summarized in Supplementary Table 1.

##### RNAPII EC•RECQ5_1-650_ complex

The movies were imported into cryoSPARC 4.02^76^ and aligned using the patch motion correction job. The CTF parameters were estimated with Patch CTF job. The images with the estimated CTF fit worse than 7Å were excluded from further data analysis. A set of 20 micrographs was randomly selected from the dataset for manual particle-picking to generate references for the template picking of the whole dataset. The overall number of 4,373,717 particles was extracted from the micrographs and binned 2x to 240 px box size (1.6672Å/px). The false positive picks and corrupted particles were removed by running multiple rounds of reference-free 2D classification resulting in 1,207,759 particles, which were used for the downstream data analysis. The *ab-initio* job was used for initial 3D classification of the data. A model corresponding to RNAPII•RECQ5_1-650_ complex, which contained 446,855 particles was further refined. Particles corresponding to RNAPII•RECQ5_1-650_ complex were re-extracted to 480 px box size (0.8336 Å/px) and another *ab-initio* job was used to generate three initial models and carry out next 3D classification of the data. *NU-Refinement* followed by the CTF refinement resulted in a 2.8 Å RNAPII-RECQ5_1-650_ map (231,041 particles). The cryo-EM density of RECQ5 and the active site of RNAPII were used to create a local mask. Two strategies were used to better characterise the active site and RECQ5 binding mechanism. In the first approach, the active site was classified first and particles belonging to the class with a well-defined active site density were selected for focused classification of RECQ5. In the second approach, the focused classification of RECQ5 was initiated, followed by the local classification of active site. The number of particles with well resolved RECQ5 and active site in the first and second approaches was 24,526 and 22,806 respectively. The final maps of RNAPII•RECQ5_1-650_ had resolutions of 3.2 Å (map 1) and 3.3 Å (map 2) (according to the FSC = 0.143 criterion). The refinement statistics are summarized in Supplementary Table 1.

##### RNAPII EC•RECQ5_1-650_ ^R502E^ complex

The movies collected using Titan Krios (TEM) were imported into cryoSPARC 4.02^76^ and aligned using the patch motion correction job. The CTF parameters were estimated with Patch CTF job. The images with the estimated CTF fit worse than 6 Å were excluded from further data analysis. A set of 20 micrographs was randomly selected from the dataset for manual particle-picking to generate references for the template picking of the whole dataset. The overall number of 787,289 particles was extracted from the micrographs and binned 2x to 200 px box size (1.6672 Å/px). The false positive picks and corrupted particles were removed by running multiple rounds of reference-free 2D classification resulting in 428,525 particles, which were used for the downstream data analysis. The *ab-initio* job was used for initial 3D classification of the data. A model corresponding to RNAPII•RECQ5_1-650_^R502E^ complex, consisting of 198,029 particles was further refined. Subsequently, the particles were re-extracted to 400 px box size (0.8336 Å/px), and another *ab-initio* job was used to generate initial model. *NU-Refinement* followed by the CTF refinement resulted in a 2.8 Å RNAPII•RECQ5_1-650_^R502E^ map. The density of RECQ5 and the active site were used to create a local mask. The RECQ5 was initially classified, and particles belonging to the class with well-defined RECQ5 density were selected for focused classification of the active site. This approach resulted in a set of 19,142 particles with well-resolved RECQ5 and active site densities but weaker nucleic acid density. The final map of RNAPII•RECQ5_1-650_^R502E^ had a resolution of 3.4 Å (according to the FSC = 0.143 criterion). The refinement statistics are summarized in Supplementary Table 1.

#### Model building and refinement

##### RNAPII EC•RECQ5_1-650_ complex

Initial PDB coordinates of the RNAPII structure were taken from PDB database (5FLM^61^). Coordinates were aligned with the density map using UCSF Chimera’s ‘Fit in Map’^77^. Subunits RPB7 and RPB4 were docked into a map with rigid body approach due to poor quality of electron density map.

Remaining subunits of *RNAPII* structure and the density map were subsequently imported into the program Coot^78^. The fitting of the coordinates into the electron density map was inspected and coordinates with no electron density were omitted from the structure. The fitting of the coordinates into the electron density map was visually inspected in coot, and if necessary, the ‘Real Space Refine Zone’ tool was employed to achieve an optimal fit of the PDB coordinates within the map. The structural refinement of the RNAPII structure was performed using the Phenix software^79^ and ISOLDE^80^. The PDB coordinates of the RECQ5 structure were taken from AlphaFold database^81^. Coordinates were fitted into the density map using UCSF Chimera’s tool ‘Fit in Map. Regions where the electron density map lacked density were omitted from the structure. The structural refinement of the RNAPII EC•RECQ5_1-650_ complex was performed using the ISOLDE. Overall model-to-map correlation coefficients were calculated using phenix.map_model_cc.

##### RNAPII EC•RECQ5_1-650_^R502E^ complex

PDB coordinates for EC-RNAPII from the EC-RNAPII•RECQ5_1-650_ complex and RECQ5, sourced from the AlphaFold database, were aligned with the density map using UCSF Chimera’s ‘Fit in Map’ tool^77^. Mutations in RECQ5 residue 502 and fitting of coordinates to the electron density map were assessed Coot, and the ‘Real Space Refine Zone’ tool was utilized when needed. Coordinates without electron in density were excluded from the structure. The structural refinement of the RNAPII structure was performed using the Phenix software^79^ and ISOLDE^80^.

### Cryo-electron tomography

#### Sample Preparation

Sample of 100 nM elongation complex (EC) of P-RNAPII was preincubated with 250 nM RECQ5^K58R^ for 15 min in RNAPII buffer. Subsequently, glutaraldehyde (0.1% final concentration) was added, and the mixture was incubated on ice for a further 15 minutes. BSA golden beads (10 nm) were concentrated 10x and transferred to RNAPII buffer. The fiducial beads were mixed with the P-RNAPII EC•RECQ5 sample at a 3:1 ratio.

The mixture (3.5 µl) was applied to freshly glow-discharged Au 300 R1.2/1.3 TEM grids (Quantifoil) coated with the monolayer of graphene and vitrified into liquid ethane using Vitrobot Mark IV (ThermoScientific) (5°C, 100% rel. humidity, 30 s waiting time, 4 s blotting time). The grids were subsequently mounted into the Autogrid cartridges and loaded to Titan Krios (ThermoScientific) transmission electron microscopes operated at 300 kV for the data acquisition.

#### Data acquisition

Tilt series were collected in SerielEM using the bidirectional (dose-symmetric) tilting scheme covering an angular range of +/−60° with 3° increment^75^. The data was collected using Bioquantum K3 direct electron detector (Ametek) operated in the counting mode at the calibrated pixel size of 1.346 Å/px and the dose rate of 15 e/px*s. Each image was acquired as a movie of 4 frames collected within 0.5 s exposure (3.6 e/Å^2^ per tilt image) resulting in the total cumulative dose for each tilt series of 146 e^−^/Å^2^. The energy selecting slit was set to 10 eV and the tilt series were collected in the defocus range of −3 to −6 µm.

#### Data processing

Pre-processing of all tilt series was performed using Warp version 1.0.9^82^. Tilt movies were corrected for whole-frame motion and aligned via patch tracking using Etomo (IMOD-version 4.11.1^83^). Tomograms were reconstructed using Warp. The deconvolution filter was used for visualization of tomograms. Warp template picker was used to identify position or RNAPII molecule in the tomograms. Cryo-EM density from single particle reconstruction of RNAPII was used as a reference for template-matching against 10 Å per pixel tomograms at a sampling rate of 7.5°. Picked particles were curated in Cube and false positive picks were removed. The initial set of subparticles (138,749 particles) was extracted to 128 px box size (4 Å/px) and subjected to several rounds of 3D classification in Relion 3.1^84^. The 3D class containing complex of RNAPII and RECQ5 from the droplets (11,866 particles) was used for further 3D refinement. The refined particles were imported into M version 1.09. Three iterations of refinement were performed starting with image-warp, particle poses and stage angles, then incorporating refinement of doming, and finally including CTF refinement of defocus. This procedure resulted in a reconstruction of the complex of RNAPII and RECQ5 at an estimated resolution 7.1 Å (according to the FSC = 0.143 criterion).

#### Model building and refinement

PDB coordinates of RNAPII EC from RNAPII EC•RECQ5_1-650_ complex and RECQ5 structure, obtained from the AlphaFold database, were aligned with the density map using UCSF Chimera’s ‘Fit in Map’ tool ^77^. Regions with missing electron density in the map were then excluded from the structure.

### Estimation of the orientations of P-RNAPII found in condensates *in vitro* and *in silico*

To assess the orientation of P-RNAPII within the condensate (either from cryo-ET data or from molecular dynamics simulations), we assumed the position of the subunits RPB4 and RPB7 (the “tower”) to be rigid with respect to the core of P-RNAPII. Using the built-in function of ChimeraX^85^ “measure center” the centers of mass of i) RPB4 and RPB7, and ii) all other RNAPII domains were obtained. This gives the respective end and start point of a vector for each individual RNAPII. The orientation of the vectors was subsequently transformed and displayed in sinusoidal (Sanson-Flasmteed) projection, displayed using Matplotlib^86^.

### Molecular dynamics (MD) simulations

In our simulations, we utilized the OpenMM molecular dynamics engine, adopting a coarse-grained approach based on the Urry hydropathy scale developed by Regy et al.^87,88^. In this model, each amino acid is represented by a single bead centered on the carbon-α atom. The model’s behavior is governed by two key parameters: µ, which modulates the strength of non-electrostatic interactions between amino acids, and Δ, which adjusts the amino acid affinity towards each other relative to the implicit solvent. We employed the optimized parameter values from Regy et al.’s publication, specifically µ = 1 and Δ = 0.08. Additionally, we incorporated parameters for phosphorylated serine from a subsequent model extension ^89^. All simulations were conducted at a constant temperature of 300 K, maintained by a Langevin Middle Integrator with a friction coefficient of 1 ps^-1^.

RECQ5 and RNAPII are both multidomain proteins with disordered and folded domains. Amino acid positions for RNAPII were taken from the existing structure (PDB: 5FLM^61^). We removed nucleic acids to prevent their interaction with surrounding proteins and the disordered the C-terminal domain (CTD) was attached to the last resolved residue of the largest RPB1 subunit of RNAPII. This disordered region includes the linker region, 52 tandem repeats of the heptapeptide consensus Y1-S2-P3-T4-S5-P6-S7, and the tail sequence of the CTD. For RECQ5, the AlphaFold predicted structure was used (Uniprot: O94762^81^). The folded domains of both proteins were kept rigid while the linkers between neighboring domains were kept as flexible chains in all simulations. Since the coarse-grained model was parameterized for treatment of disordered regions, keeping folded regions rigid ensures that the protein does not unfold during the simulation. The regions that were kept rigid are shown in Supplementary Table 5. The net charge per residue (NCPR) was calculated using localCIDER, a tool-box for sequence analysis developed by Alex Holehouse as a part of the Pappu Lab.

In our simulations, we considered the reported low micromolar binding affinity of RECQ5’s SRI domain to P-CTD as a reference point^90^. However, the SRI region of RECQ5 was kept rigid in our model, which could potentially inhibit this interaction. To approximate the simulated affinity between P-CTD and RECQ5’s SRI domain, we introduced an additional potential. Specifically, we focused on the five SRI residues (V920, F938, K939, R943, H947), known to bind P-CTD, by adding positive charges to the five residues that could only interact with phosphoserines^90^. We employed umbrella sampling to estimate the dissociation constant, using the center of mass separation distance as the reaction coordinate. A harmonic bond with a force constant of 2 kcal/nm was applied to maintain this separation distance. The range of separation distances spanned from 6 to 1 nm, using 30 sampling windows to cover this range. Each window was simulated for 100 ns, recording data every 0.1 ns, to thoroughly sample the interaction dynamics between the two molecules. The simulation was then reversed (separation distance of 1 to 6 nm) to ensure there was no hysteresis in the measurement. At a center of mass separation distance of 3.5 nm, the minimal distance between the SRI and diheptad of CTD exceeds the cutoff from all pair potentials. Therefore, the free energy at 6 nm is well within the unbound state. The weighted histogram analysis method (WHAM) was used to generate the free energy landscape, which was integrated to determine the binding affinity between the two molecules^91^. A table of the phosphorylation states tested, along with the measured K_D_ can be found in Supplementary Table 6.

The radius of gyration (Rg) for the disordered regions of each protein, RECQ5_IDR_, and the CTD of RNAPII, was calculated every 1 ns for the last 4 µs of a 5 µs simulation run.

We verified that the choice of parameterization accurately reproduces experimentally observed condensate formation of the following individual species: RECQ5_IDR_, CTD, and P-CTD. For each protein, 100 copies were placed in a 15 x 15 x 150 nm box. The proteins were placed parallel to each other, in a straight line along the Z-axis, in the center of the box. This system was then simulated in NVT ensemble at 300 K for 5 µs. To analyze the trajectory, the simulation was centered, ensuring the center of the largest protein cluster was in the box center. The density was then fit to a hyperbolic tangent function, averaged over every 2 ns for the last 1 µs of each simulation. This procedure has been well documented and shared by Tesei & Larsen ^92^

The condensate forming ability of RECQ5 was demonstrated by placing 70 copies of RECQ5 in a cubic 60 x 60 x 60 nm box and simulated for 10 µs using an NVT ensemble. Since there is no compression involved, any condensate formation would be fully spontaneous. By studying spontaneous RECQ5 condensate formation we avoid defects that may be present due to the large, folded regions in the slab model. This may come in the form of non-uniform density throughout the slab, which may cause unforeseen problems, and was therefore avoided entirely. The inter-protein contacts are defined as any amino acids with a separation distance less than 1.5 nm.

In simulations where both P-RNAPII and RECQ5 were present, the RECQ5_KIX_ domain was bound to the RPB1 jaw, a condition corroborated by our cryo-ET density analyses. To maintain this interaction, we introduced a bond between residue D529 of RECQ5 and residue V1247 of RNAPII. The amino acid positions for the KIX domain were adjusted to match those observed in the cryo-ET densities. During simulation setup, each P-RNAPII molecule was inserted with one RECQ5 molecule bound via the IRI domain.

Condensates composed of RECQ5 and P-RNAPII were created by randomly placing copies of each protein into a box of 700 x 700 x 700 nm while avoiding protein overlaps. After insertion of all proteins, the box was compressed isotopically in an NPT simulation for 0.1 µs at 1 bar and 150 K, using a Monte Carlo barostat. The final size of the box was then roughly 230 nm in all dimensions for systems with 10 copies of P-RNAPII, and 420 - 600 nm for larger systems with 30, 50 and 60 copies of P-RNAPII. The compressed system was then moved into the original box (700 x 700 x 700 nm) and equilibrated using a NVT ensemble. Equilibration was determined using an autocorrelation function, to determine when the valency of the RECQ5_IDR_ domain, P-CTD domain, and center of mass separation between P-RNAPII were no longer dependent on the starting configuration. The equation used for autocorrelation can be seen in eq S1, and the cut-off of 1/e was chosen to signify the end of equilibration phase ^93^. In eq S1 & 2, CA is the autocorrelation function, A is the measurable at time t, and N is the number of timesteps (100 per microsecond).

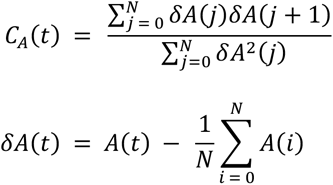

Four concentrations of RECQ5 were studied, using 10 copies of P-RNAPII, to infer the relative ratio seen *in vitro*, which would then be used in larger systems. The ratio was chosen by comparing the average separation distance between the center of mass of RNAPII cores, to that seen experimentally. The ratios (P-RNAPII•RECQ5) studied were 1:5, 1:7, 1:8, 1:9, and the RNAPII separation distance was measured after the condensates had equilibrated.

Finally, systems with 30, 50 and 60 copies of P-RNAPII, were built using the same method described above for 10 P-RNAPIIs. Seven copies of RECQ5 were inserted for each P-RNAPII, as informed by the previous section. Each condensate was analyzed to determine the network connectivity. This was determined by measuring the valency of interacting proteins. The proteins were considered to be interacting, when the distance between interacting pairs (RECQ5_IDR_•RECQ5_IDR_ or SRI•CTD) were closer than 3 nm. The valency for these contacts, were measured over 5 µs in 100 ns intervals after the condensates had equilibrated.

## Data and materials availability

Cryo-EM/ET maps are deposited in the Electron Microscopy Data Bank under accession numbers EMD-18573 (the RNAPII EC•RECQ5_1–650_ complex), EMD-18683 (the RNAPII EC•RECQ5_1–650_^R502E^ complex), EMD-18685 (RNAPII EC•RECQ5^ΔIDR^), and EMD-18682 (the RNAPII EC•RECQ5 complex from subtomogram averaging). The atomic models are deposited in the Protein Data Bank under accession numbers 8QQ2 (the RNAPII EC•RECQ5_1–650_ complex), 8QW9 (the RNAPII EC•RECQ5_1–650_^R502E^ complex), and 8QW8 (the RNAPII EC•RECQ5 complex from subtomogram averaging). Other structures used in this study are available under PDB accession numbers 5LB3, 8A40, and 6TED. Small angle X-ray scattering data were deposited into Small Angle Scattering Biological Data Bank under the following accession numbers: SASDTK7, SASDTL7 (RECQ5 wild-type) and SASDTM7, SASDTN7, SASDTP7, SASDTQ7, SASDTR7 (RECQ5_625-825_). Code, simulation data, and basic parameter files used in the MD work can be found at https://github.com/shakespearemorton/RNAP_RECQ5_Publication/, and any further data will be made available upon reasonable request. Requests for reagents, plasmids, and cell lines used in this study should be directed to the corresponding authors.

## Acknowledgments

We thank Dr. Florian Schur and Julia Datler for discussions and preliminary processing of the cryo-ET data. We thank Dr. Pavel Janscak for providing plasmids for purification of wild-type RECQ5 from *E. coli*. We thank Dr. Susanne Kassube for providing the pGEMTerm plasmid for the transcriptional read-through experiment. We thank Dr. Katerina Sedova and Matyas Lanicek for their assistance with the experiments. The main funding was provided by the Czech Science Foundation to M.S. (grant no. 20-21581Y), by the MEYS CR to R.S. (grant no. CZ.02.01.01/00/22_008/0004575 RNA for therapy), the project National Institute of virology and bacteriology to R.V. (Programme EXCELES, LX22NPO5103) - Funded by the European Union - Next Generation EU. Additional funding was provided by the Czech Science Foundation to R.S. (grant no. 21-24460S) and to V.B. (grant no. 22-25365S) and the European Research Council to R.S. (grant no. 649030) and to R.V. (grant no. 101001470), and by the project National Institute for Cancer Research to V.B. (Programme EXCELES, LX22NPO5102 - European Union - Next Generation EU. We acknowledge BIC, CRYO, NMR, Proteomics of CIISB, Instruct-CZ Centre and the core facility CELLIM of CEITEC supported by MEYS CR (grant no. LM2018127) and (grant no. LM2018129 Czech-BioImaging), respectively. Computational resources were provided by the CESNET, CERIT Scientific Cloud, and IT4 Innovations National Supercomputing Center by MEYS CR through the e-INFRA CZ (ID:90254).

## Author contributions

MS and RS conceptualized and organized the study. MS, RV, and RS wrote the manuscript, with input from all authors. MS, JN, VB, RV, and RS supervised the work, contributed to the data interpretation, and provided financial resources. MS, KS, JN, KK, RV, and RS designed experiments and methodology. MS, KS, WSM, VK, KL, MK, JN, KK, and RS performed experiments, data analysis, and visualization.

## Extended Data Figures

**Extended Data Fig. 1:**
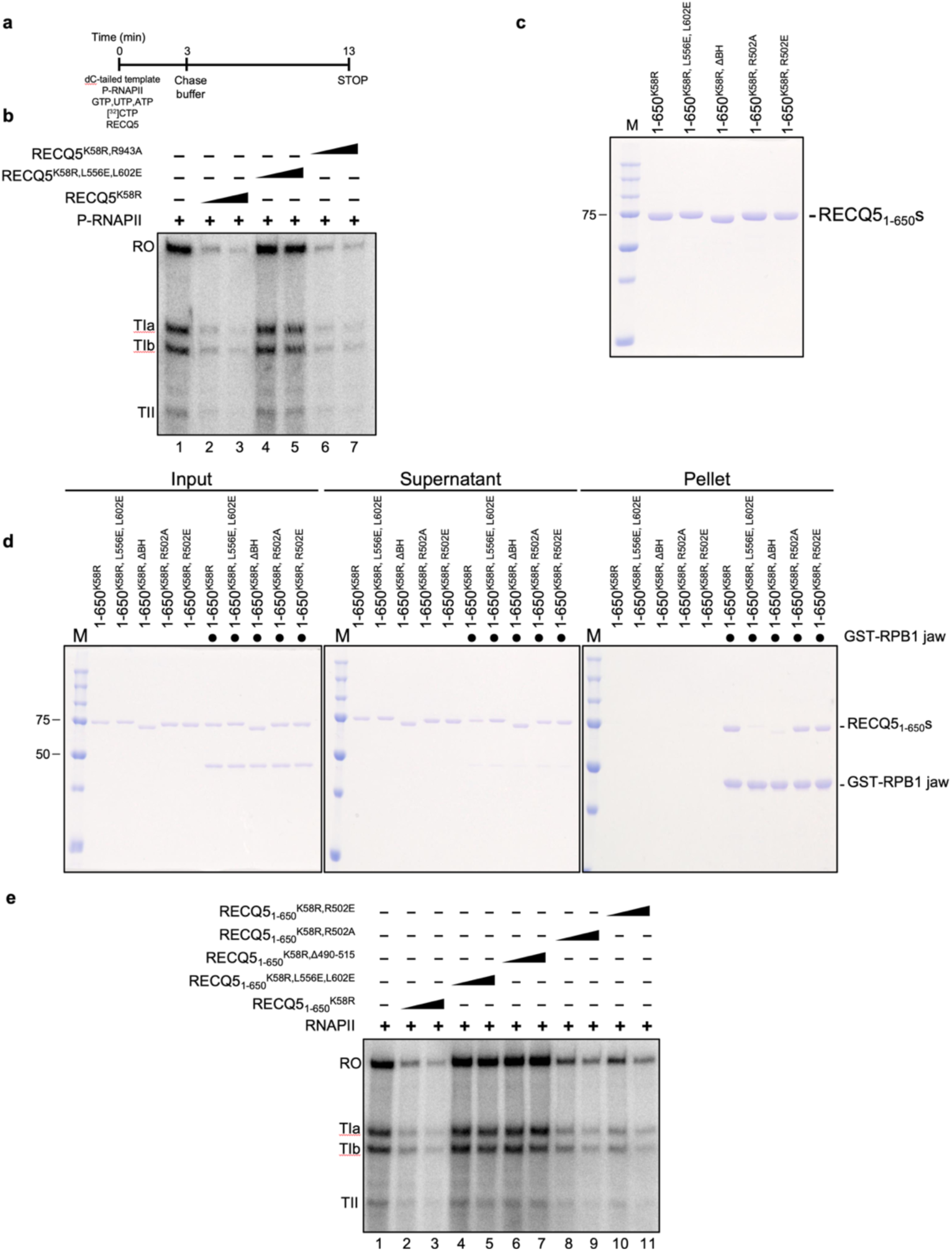
**a**, Schematic overview of the *in vitro* transcription assay. The entire reaction was assembled on ice, started by the addition of α-[^32^P]-CTP, followed by 3 min incubation at 30°C, thereby uniformly labeling the nascent RNA. Subsequently, excess of unlabeled CTP was added, and the reaction was continued for additional 10 min. After which, the reaction was stopped with proteinase K and fractioned on denaturing PAGE gel. **b**, Effect of RECQ5 substitutions on the activity of phosphorylated RNAPII (P-RNAPII). The same experiment as in Fig. 1a; the entire gel, showing the pausing products (TIa, TIb, and TII), as well as the run-off products is depicted. **c**, Purified variants of RECQ5_1-650_. Scan of an SDS-PAGE gel showing the indicated RECQ5_1-650_ variants. BH: Break Helix (RECQ5 AA 495-515). **d**, Determination of the binding properties of the RECQ5_1-650_ variants towards RPB1 jaw domain. A scan of SDS-PAGE gels from *in vitro* pull-down experiment. The same experiment is shown in Fig. 1d; the entire reaction is depicted. **e**, The brake helix and R502 are required for transcription repression. Transcription assay as in a, performed with RNAPII. The same experiment as in Fig. 1e; the entire gel, showing the pausing products (TIa, TIb, and TII), as well as the run-off products is depicted.

**Extended Data Fig. 2:**
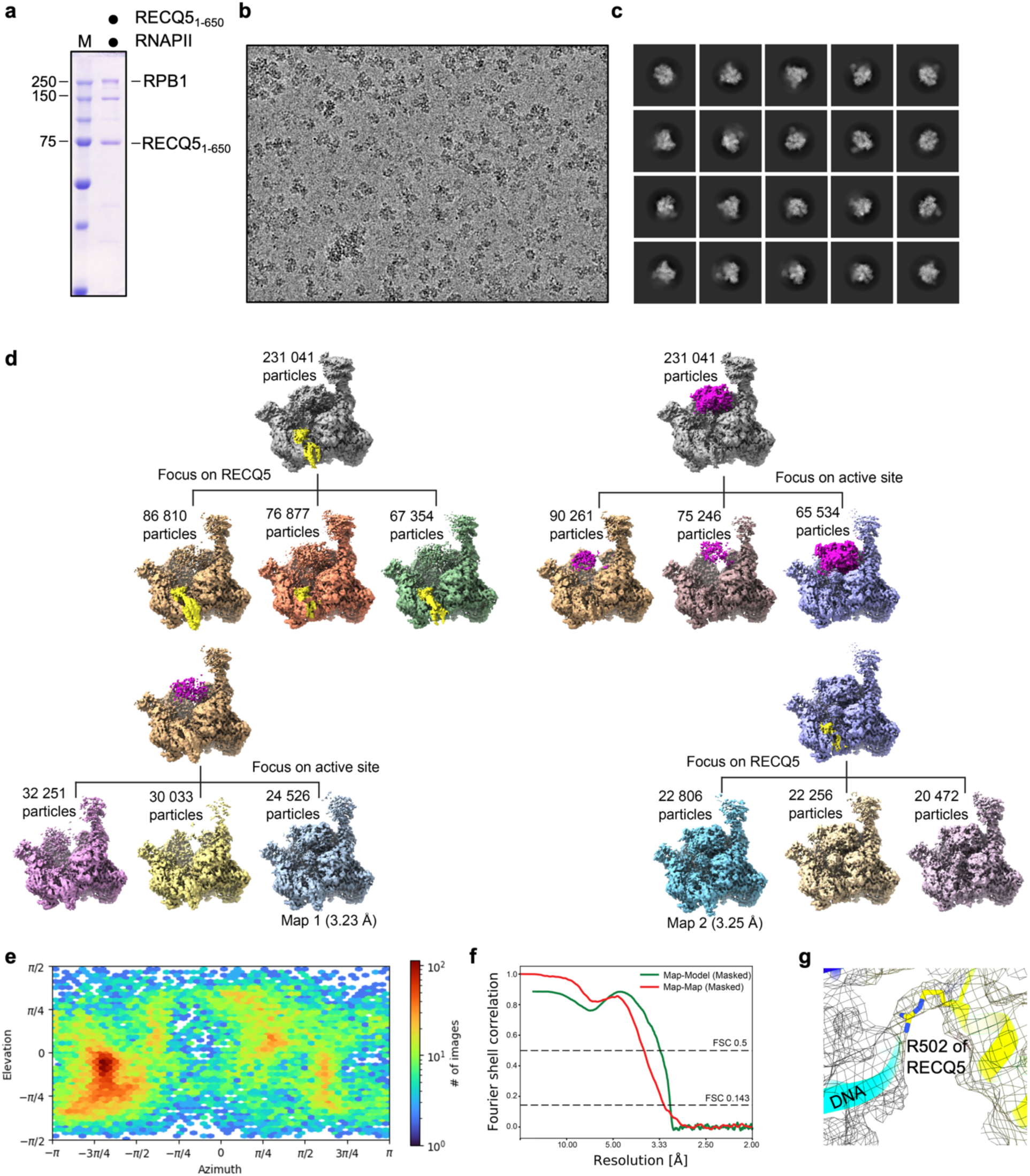
Data acquisition, processing, and data quality metrics for the RNAPII EC•RECQ5_1-650_ complex. **a**, SDS-PAGE of the purified complex. **b**, Representative denoised micrograph. **c**, 2D classes. **d**, Sorting and classification tree of the complex. Two independent sorting pathways yielded essentially identical maps (Map 1 and Map2). **e**, Heat map for distribution of particles for the final 3D reconstruction. **f**, FSC curves of overall model (Map1). **g**, Close-up of the cryo-EM map (Map 2) for the interaction of the brake helix with DNA.

**Extended Data Fig. 3:**
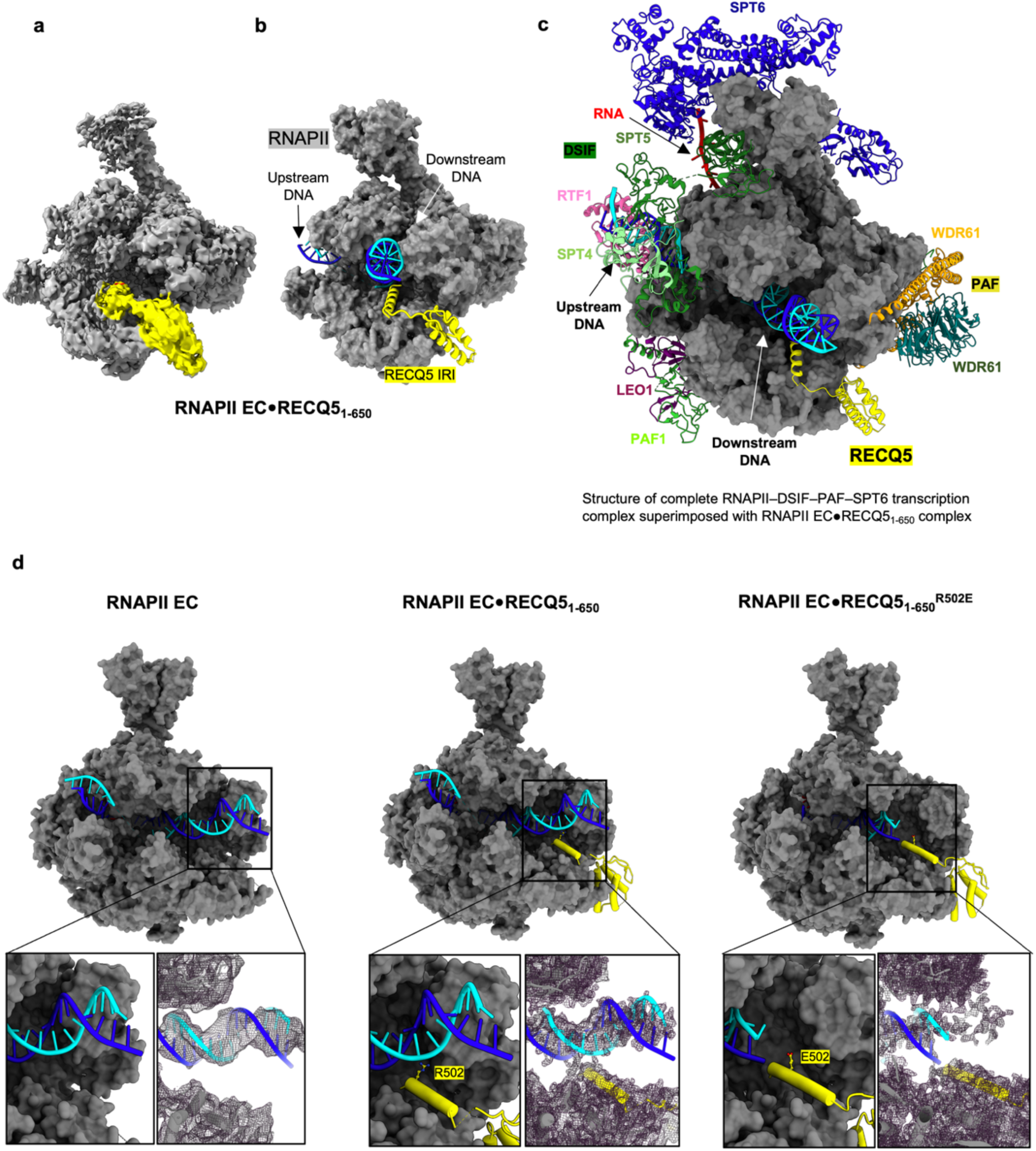
Comparison of cryo-EM structures. Final cryo-EM map (**a**) and structure (**b**) of the RNAPII EC•RECQ5_1-650_ complex. **c**, Overlay of the RNAPII–DSIF–PAF–SPT6 elongation complex (PDB ID: 6TED) with the RNAPII EC•RECQ5_1-650_ complex. **d**, Comparison of the cryo-EM maps and structures of the free RNAPII EC and bound to RECQ5_1-650_ and RECQ5_1-650_^R502E^. The comparison highlights the downstream DNA and its recognition by the brake helix of RECQ5. The model structure of RNAPII EC in the apo form was build using the coordinates from the RNAPII EC•RECQ5_1-650_ complex.

**Extended Data Fig. 4:**
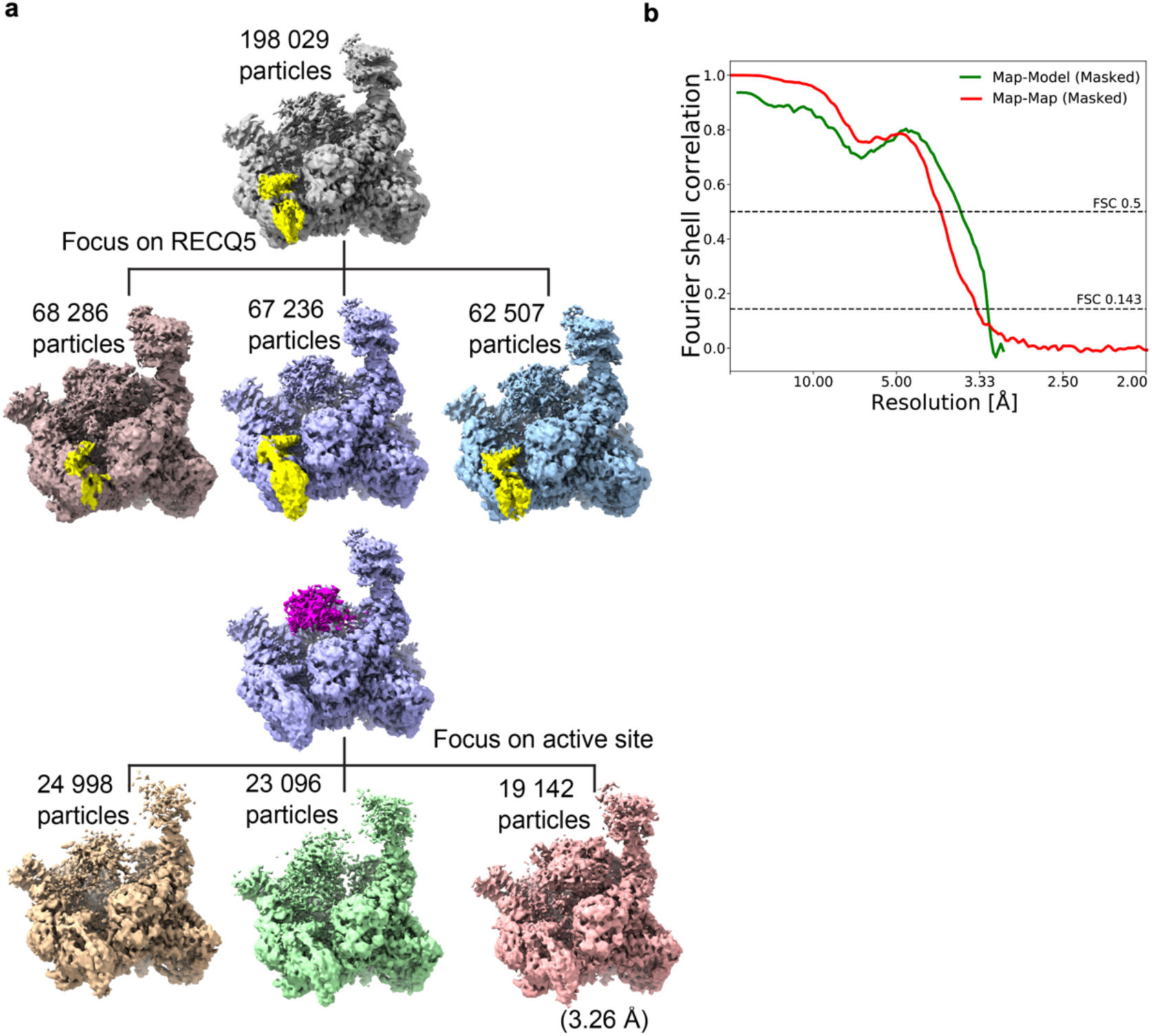
**a**, Processing and data quality metrics for the RNAPII EC•RECQ5_1-650_^R502E^ complex. Sorting and classification tree of the complex. **b**, FSC curves of overall model.

**Extended Data Fig. 5:**
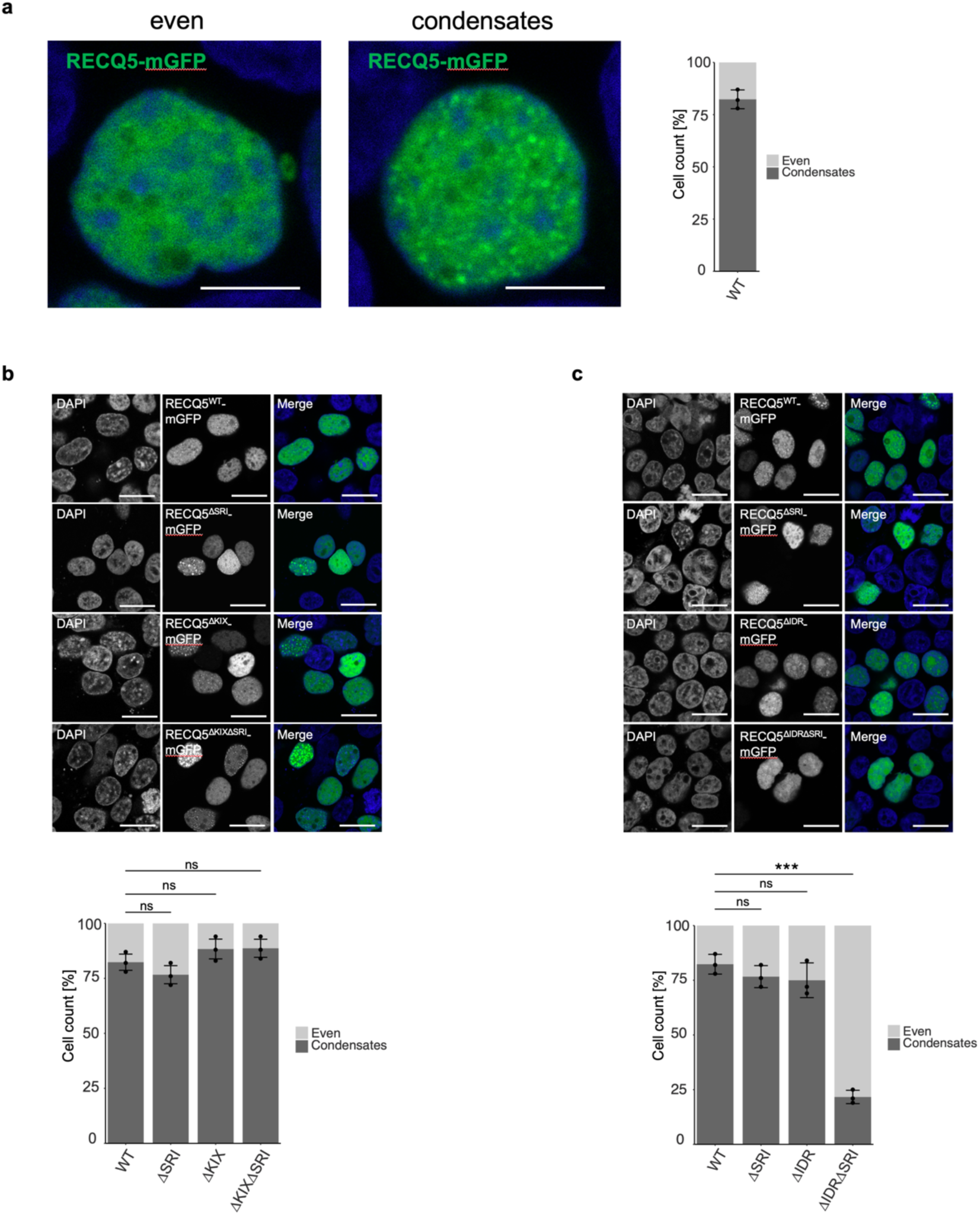
Ectopically expressed RECQ5 forms condesates *in vivo*, which require intact IDR and SRI domains. **a**, A representative micrographs of HEK293T nuclei, ectopically expressing RECQ5-mGFP, in which RECQ5 is either evenly distributed (left) or form condesates (middle). Pictures are depicted as merge of DAPI signal (staining for DNA) and mGFP. Scale bar represents 5 µm. **b**, KIX and SRI domains, responsible for interaction with the RNAPII, do not affect the ability of RECQ5 to form condensates *in vivo*. Representative micrographs of cells stained for DAPI (nuclei), expressing RECQ5-mGFP, RECQ5^ΔSRI^-mGFP, RECQ5^ΔKIX^-mGFP, and RECQ5^ΔKIXDΔSRI^-mGFP (as indicated on the figures), and merge of both channels. Scale bar represents 10 µm. **c**, Simultaneous inactivation of IDR and SRI domains is required to abolish the ability of RECQ5 to form condensates *in vivo*. Representative micrographs of cells stained for DAPI (nuclei), expressing RECQ5-mGFP, RECQ5^ΔSRI^-mGFP, RECQ5^ΔIDR^-mGFP, and RECQ5^ΔIDRΔSRI^-mGFP (as indicated on the figures), and merge of both channels. Scale bar in B and C represents 10 µm. The bar charts in the figure represent mean of cells (n=100) whose nuclei contain RECQ5-mGFP in condensates (dark grey). The error bars represent standard deviation from n=3 experiments. Statistical significance was determined by unpaired t-test.

**Extended Data Fig. 6:**
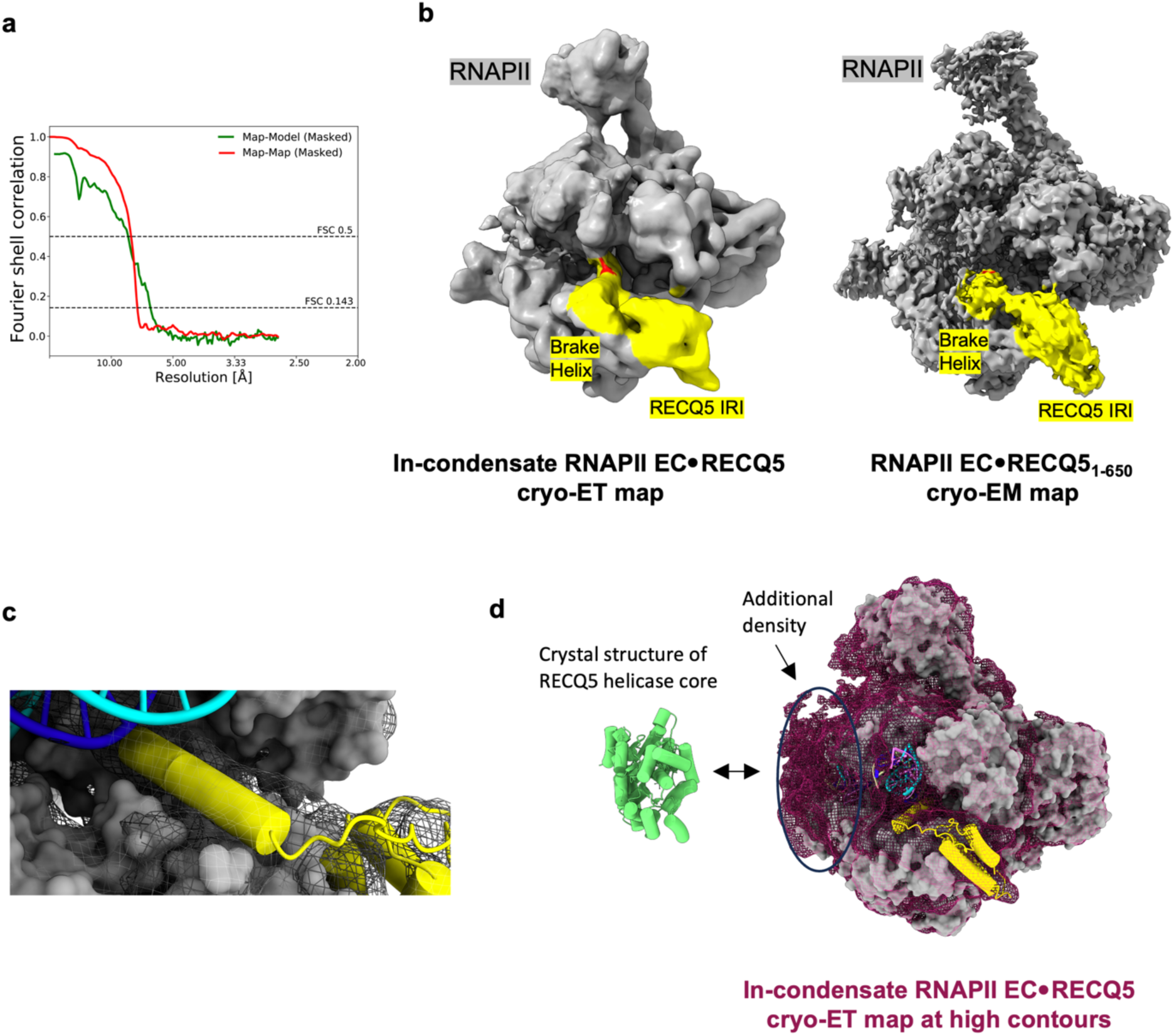
Analysis of the cryo-ET map of the RNAPII EC•RECQ5 complex. **a**, Map-to-map and map-to-model FSC curves. **b**, Comparison of the cryo-ET map of the RNAPII EC•RECQ5 complex with the cryo-EM map of the RNAPII EC•RECQ5_1-650_ complex. **c**, Close-up view of the extended brake helix of RECQ5 (in yellow) with cryo-ET map, shown as black mesh. **d**, Overall view of the cryo-ET map of the RNAPII EC•RECQ5 complex at high contours in bordeaux mesh, with RNAPII as grey surface, RECQ5 IRI as yellow cartoon, DNA as cyan-blue cartoon. Crystal structure of RECQ5 helicase core (PDB ID: 5LB3), shown as green cartoon, has a similar size as the additional density in cryo-ET map.

**Extended Data Fig. 7:**
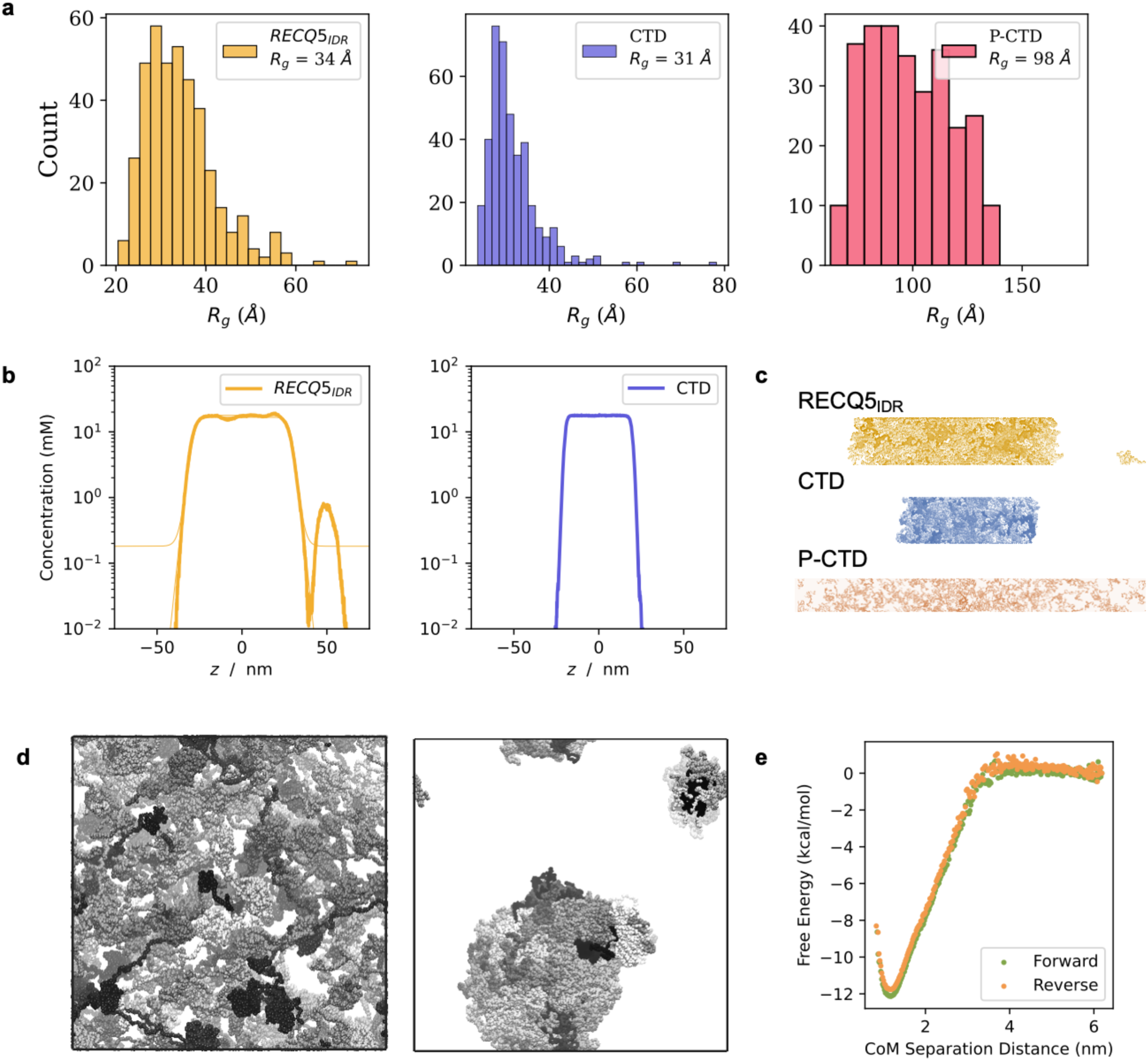
**a**, A histogram of the radius of gyration (R_g_) for RECQ5_IDR_, CTD, and P-CTD (Y1-pS2-P3-T4-pS5-P6-S7). The RECQ5_IDR_ is within the range determined from this work by SAXS (34-39 Å), and the CTD expands upon phosphorylation. **b**, Density profiles of the simulated box with RECQ5_IDR_, CTD, and S5P-CTD along the z-axis. The graph exhibits the local density of molecules within a defined volume. The P-CTD displays a broad distribution roughly equal throughout the box, reflecting a gas-like distribution.). **c**, Comparative visualization of the final molecular arrangements in simulations for RECQ5_IDR_, CTD, and P-CTD. Condensates from RECQ5_IDR_ and CTD formed a slab due to the applied periodic boundary conditions. P-CTD resulted in a dispersed solution. **d**, Seventy copies of REQC5 in a 60 x 60 x 60 nm periodic box at the beginning of the simulation (Left) and after 17 μs (Right). Two condensates have formed spontaneously within the simulated timescale, demonstrating RECQ5’s ability to spontaneously form condensates. Each RECQ5 is given a different color to help distinguish their positions. **e**, Free energy profiles for the interaction between the phosphorylated diheptad of CTD (Y1-pS2-P3-T4-pS5-P6-S7) and the SRI domain of RECQ5. The distance between proteins’ center of mass (CoM) was systematically reduced from 5 nm to 1 nm and then reversed, confirming the robustness of the sampling through the absence of hysteresis. The binding affinity in Mol is shown for each of the charges placed on the SRI domain.

**Extended Data Fig. 8:**
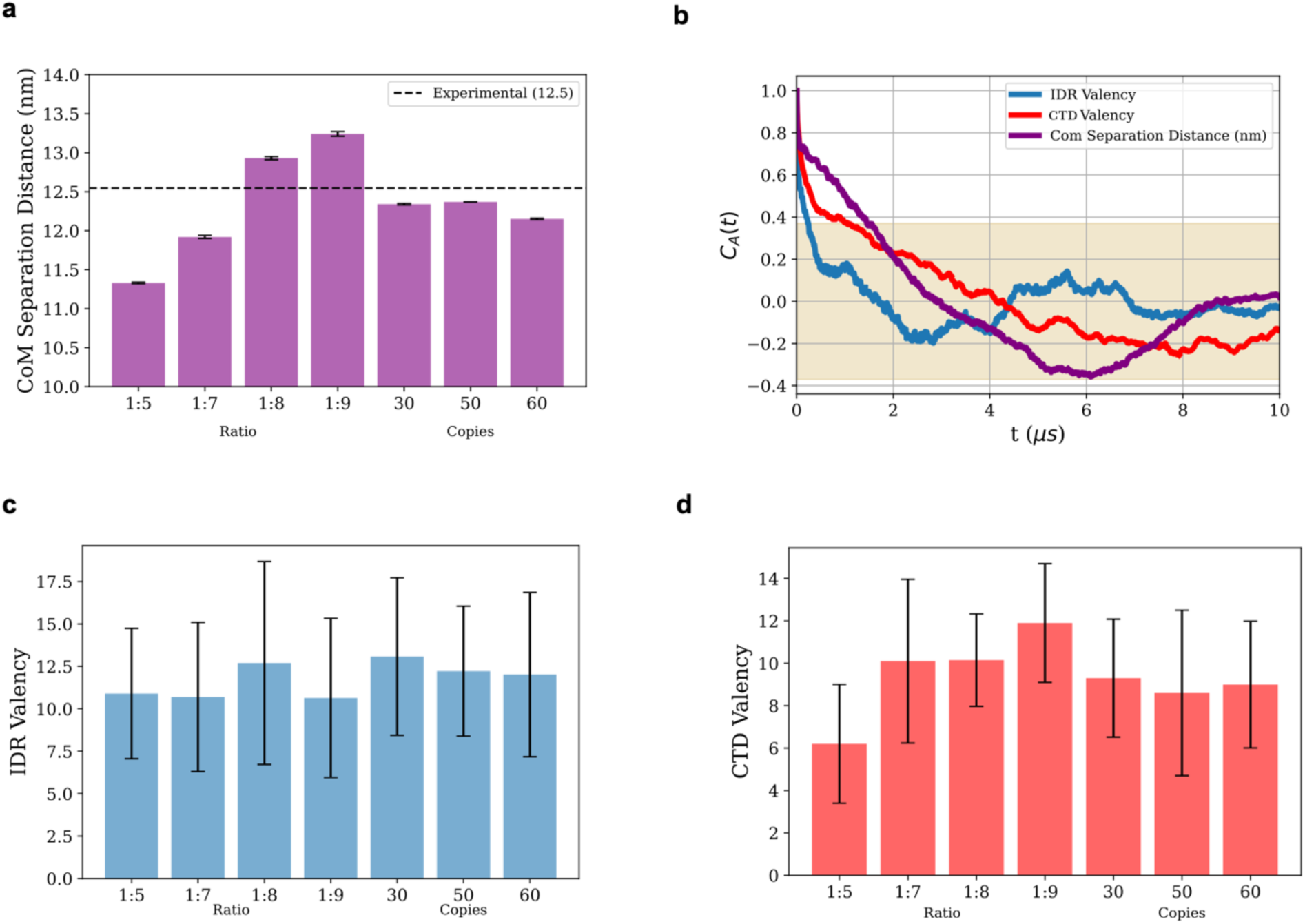
**a**, The center of mass (CoM) separation distance between P-RNAPII within condensates containing 10 copies of P-RNAPII and 50, 70, 80, or 90 copies of RECQ5. The CoM separation distance is also shown for condensates with 30, 50, and 60 copies of P-RNAPII, and 7 RECQ5 for each P-RNAPII. The black dotted line indicates the separation distance seen experimentally. **b**, Autocorrelation function of three observables: IDR valency, CTD valency, and CoM separation distance between P-RNAPII. When all three values are less than 1/e (∼ 0.37) away from 0, the condensate is considered equilibrated. This is an example from the first 10 microseconds of the simulation with 10 P-RNAPII and 50 RECQ5. **c**, Average valency between RECQ5_IDRs_ in condensates with varying ratios (of P-RNAPII:RECQ5 with 10 copies of P-RNAPII) and with increasing size (30, 50 and 60 copies of P-RNAPII) using a 1:7 ratio. Error-bars represent one standard deviation from the mean valency. **d**, Average of binding between SRI and P-CTD in condensates with varying ratios (of P-RNAPII:RECQ5 with 10 copies of P-RNAPII) and number of P-RNAPII copies (condensates with 30, 50 or 60 P-RNAPIIs) using a 1:7 ratio. Error-bars represent one standard deviation from the mean valency.

